# Developmental temporal programs govern and maintain neuronal identity and function in adult circuits

**DOI:** 10.64898/2026.01.28.702037

**Authors:** Adil Rashid Wani, Rahail Ashraf, Riley Woerner, Zoe Marriner, Michael Velasquez, Tulaib Azam, Qussin Joo, Monica Dus, Mubarak Hussain Syed

**Affiliations:** Neural Diversity Lab, Department of Biology, University of New Mexico, 219 Yale Blvd NE, Albuquerque, NM 87131, USA; Department of Molecular, Cellular and Developmental Biology, College of Literature, Science, and the Arts, The University of Michigan, Ann Arbor, MI 49109, USA

## Abstract

Developmental transcriptional programs establish neuronal diversity and circuit assembly, but how these regulators continue to shape and preserve neuronal identity remains unresolved. The *Drosophila* central complex comprises precisely wired circuits of diverse neuronal types that coordinate complex behaviors. Within this structure, dorsal fan-shaped body (dFB) neurons integrate internal-state signals to regulate sleep, feeding, and energy homeostasis, yet how distinct dFB subtypes are specified and maintained remains unknown. More broadly, whether developmental transcription factors continue to act in mature neurons to preserve neuronal identity is poorly understood.

Here, we define the developmental origin and molecular regulation of neuronal identity in 84C10-labelled dFB neurons that innervate layers 6-7 of the FB and contribute to nutrient sensing and metabolic adaptation. Using lineage tracing, clonal analysis, and birth dating, we show that dFB neurons are generated late in development and arise from two distinct type II neural stem cell lineages, dorsolateral 1 and dorsomedial 4. These dFB neurons continue to express the late temporal transcription factor, ecdysone-induced protein 93 (E93), in adulthood. Post-mitotic depletion of E93 results in progressive neuronal loss and ectopic expansion of axonal arborizations across FB layers and, notably, a marked reduction in vesicular glutamate transporter (vGLUT) expression. These defects are accompanied by impaired increases in fat-to-lean mass ratios in response to high-sugar feeding.

Together, our findings identify E93 as a post-mitotically retained temporal transcription factor that maintains neuronal survival, laminar connectivity, and neurotransmitter identity, revealing that developmental temporal programs are redeployed in adulthood to sustain neuronal identity and function.

## Introduction

Complex behaviors arise from neural circuits composed of diverse neuron types whose identities are specified during development and maintained throughout life[1,2]. These neuron types differ in morphology, connectivity, molecular identity, and function, enabling the construction of circuits that support innate and adaptive behaviors[3,4]. How transcriptional programs that operate in neural stem cells (NSCs) are translated into stable, functionally relevant identities in mature neurons remains a central question in neuroscience[5,6].

Across the nervous system, neuronal diversity is generated through the integration of multiple developmental axes, including spatial patterning of neural progenitors[7,8], temporal progression of progenitor competence[5,9], and binary fate decisions at neuronal birth[10,11]. Temporal transcription factors (TTFs), expressed sequentially in neural stem cells, are well established as key regulators of neuronal fate specification during neurogenesis[9,12,13]. These temporal programs define when specific neuronal classes are produced and contribute to their initial molecular and anatomical identities[5,14]. However, whether and how such temporal programs persist beyond neurogenesis to maintain postmitotic neuronal identity, connectivity, and physiological function remains poorly understood[6,15].

The *Drosophila melanogaster* central complex (CX) provides a powerful system for addressing this problem[16,17]. The CX is a conserved brain center across insects that is required for innate behaviors, including locomotion, navigation, feeding, and sleep[18–20]. It comprises hundreds of anatomically and functionally distinct neuron types, organized into stereotyped neuropil[21,22]. Many CX neurons are generated by type II NSCs, a small set of NSCs that undergo prolonged temporal patterning during larval development and generate large, diverse neuronal lineages[23–26]. These neurons innervate distinct CX neuropils and form layered and columnar circuit architectures[22].

Within the CX, neurons projecting to the dorsal fan-shaped body (dFB) play key roles in regulating behavioral state and metabolic homeostasis[27],[28]. Previous work identified a population of dFB neurons labeled by the 23E10 driver that are specified during a late temporal window by the late TTF, ecdysone-induced protein 93 (E93) and are required for sleep regulation[27,29]. A related population labeled by the 84C10 driver innervates overlapping layers of the dFB and has been implicated in feeding-related behaviors and nutrient homeostasis[30–32]. Despite their functional relevance, the developmental origin, molecular identity, and regulatory logic underlying the specification and maintenance of 84C10 dFB neurons remain largely unexplored.

Here, we use 84C10 dFB neurons as a model to investigate how late temporal programs shape neuronal identity and function. We show that 84C10 dFB neurons are generated during a late temporal window from two type II NSC lineages, and that this population exhibits molecular heterogeneity. We identify the late temporal transcription factor E93 as a key regulator of 84C10 dFB neurons, acting not only during neurogenesis but also post-mitotically to maintain neuronal identity and to regulate precise axonal targeting and neurotransmitter expression. Disruption of E93 in 84C10 dFB neurons compromises neuronal survival, alters circuit architecture, and impairs metabolic adaptation to dietary challenge.

Together, our findings demonstrate that TTFs can function beyond progenitor stages to actively maintain adult neuronal identity and physiological function, providing a developmental framework for understanding how complex innate behavior circuits are built and sustained.

## Results

### The dFB neurons represent a heterogeneous neuronal population

The dorsal fan-shaped body (dFB) regulates sleep, feeding, and metabolic state[27,28,33]. Although it has often been treated as a single functional population, recent EM reconstructions and lineage analyses reveal that the dFB comprises multiple tangential neuron subtypes that differ in developmental origin, connectivity, and molecular identity[22,24,29,34,35],[36]. These findings raise a fundamental developmental question: how does this cellular and circuit-level heterogeneity arise? Specifically, what lineage programs generate distinct dFB subtypes with divergent wiring and functional roles? Thus, rather than viewing the dFB as a uniform functional node, we treat it as a heterogeneous circuit module whose diversity must be explained by its developmental logic.

To begin addressing this question, we first sought to define with greater precision the cellular composition of commonly used genetic access lines that target dFB neurons. Much of what is known about dFB function derives from studies using the 23E10 driver, and our previous work demonstrated that this population arises from type II neuroblasts and plays a central role in sleep regulation[29]. However, growing anatomical and molecular evidence indicates that 23E10 labels a heterogeneous population of dFB neurons[36], raising the possibility that functionally distinct subtypes are embedded within this broadly defined group.

The 84C10 driver also labels long-field tangential neurons with somata in the protocerebral posterior lateral (PPL) region and axonal projections to layers 6-7 of the FB. Given the emerging view of dFB heterogeneity, a critical first question was whether 84C10 identifies the same neurons labeled by 23E10 or instead marks an overlapping but developmentally distinct subset. Clarifying the relationship between these two widely used drivers is essential for interpreting prior functional studies and for dissecting subtype-specific developmental programs.

To directly compare the two driver lines, we used intersectional genetics to simultaneously express UAS-RFP under 23E10-GAL4 and LexAop-GFP under 84C10-LexA in the same animals. In adult brains, approximately eight neurons co-expressed GFP and RFP (pseudo color magenta), indicating a shared population labeled by both drivers (Figs 1A and 1D). In addition, several neurons (∼7) were labeled exclusively by either 23E10 (RFP) (pseudo color magenta) or 84C10 (GFP), demonstrating that while the overlap between the two lines is substantial, it is incomplete (Figs 1B and 1C). This partial overlap is consistent with previous reports showing that the 23E10 driver encompasses multiple molecularly and anatomically distinct dFB neuron subtypes[36]. Because this comparison relied on two independent binary expression systems (GAL4 and LexA), we next validated the shared population using an expression-independent intersectional split-GAL4 approach.

**Fig 1.**
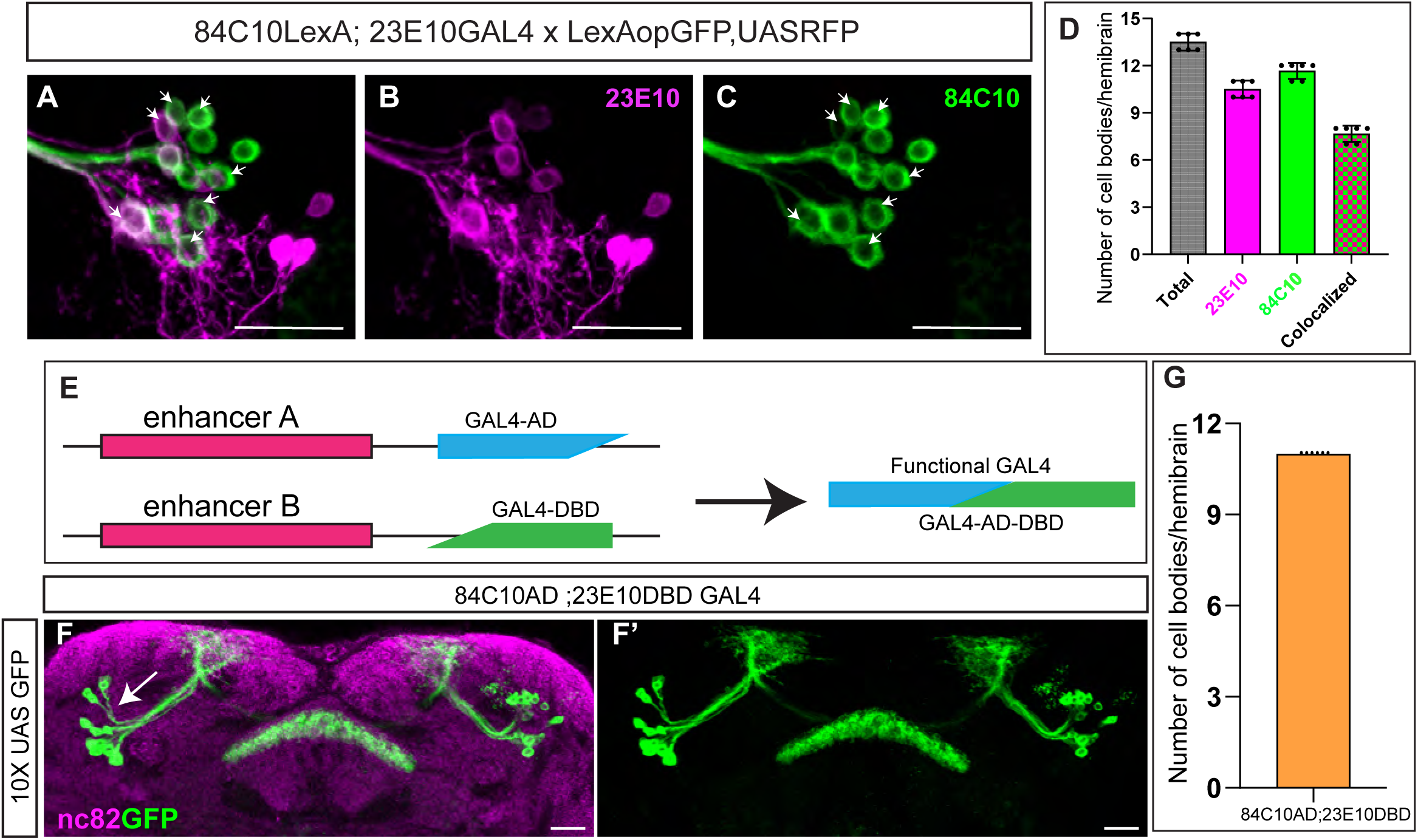
Intersectional labeling reveals partial overlap between 84C10 and 23E10 dFB neurons. (A) Dual labeling of dFB neurons using the LexA–LexAop system (84C10LexA > LexAop-GFP) and the GAL4–UAS system (23E10-GAL4 > UAS-RFP) (pseudo color magenta) reveals partial overlap between the two driver lines. Approximately eight neurons co-express GFP and RFP, indicating a shared population. White arrows indicate co-labeled cell bodies. (B) Representative confocal image showing dFB neuron cell bodies labeled by 23E10-GAL4 > UAS-RFP (pseudo color magenta). (C) Representative confocal image showing dFB neuron cell bodies labeled by 84C10LexA > LexAop-GFP. (D) Quantification of dFB neurons labeled by 23E10-GAL4, 84C10LexA, and those co-labeled by both drivers using the dual-system approach. Bars represent mean ± SD. (E) Schematic illustrating the Split-GAL4 strategy. The activation domain (AD) from 84C10 and the DNA-binding domain (DBD) from 23E10 reconstitute functional GAL4 only in neurons where both enhancers are active. (F, F′) Representative confocal images of dFB neurons labeled by the 84C10-AD;23E10-DBD Split-GAL4 line crossed to UAS-GFP. This intersectional strategy selectively labels neurons common to both 84C10 and 23E10 driver lines. GFP-positive neurons are shown in green (maximum projection), and neuropil is labeled with nc82 (magenta). Images are shown as FB-focused projections. (G) Quantification of dFB neurons labeled by the 84C10-AD;23E10-DBD Split-GAL4 line. An average of ∼11 neurons per hemibrain are labeled, corresponding to the shared population identified in the dual-system experiments. Scale bars, 20 µm, *n* = 6 adult hemibrains. Error bars represent mean ± SD. White arrows indicate neuronal cell bodies.

We generated a split-GAL4 line by combining 84C10-AD with 23E10-DBD, enabling GAL4 reconstitution only in cells in which both enhancer elements are active (Figure 1E). When crossed to UAS-GFP, this intersectional line labeled a discrete set of ∼11 neurons within the adult dFB (Figs 1F, 1F′, and 1G), which closely corresponds to the shared population of neurons identified in the dual-system experiments (Fig 1D).

Together, these findings establish that 84C10 labels a major subset of the canonical 23E10-defined dFB neurons. Although the two drivers show broad overlap, each also marks additional neurons, indicating that 23E10 and 84C10 capture overlapping but non-identical components of the dFB population.

### 84C10 dFB neurons derive from type II NSCs

The *Drosophila* CX comprises approximately 3,000 neurons representing ∼250 distinct neuronal types[37]. A substantial fraction of both neurons and glia in the adult CX is derived from 16 type II NSCs, which generate diverse lineages through intermediate neural progenitors[5,24],[38–41]. The 84C10-labeled dFB neurons constitute a small, genetically accessible population that has been implicated in feeding-related behaviors[31,32]. Given their functional importance, we next sought to determine their developmental origin.

To test whether 84C10 dFB neurons arise from type II NSCs, we employed an intersectional genetic strategy that restricts reporter expression specifically to the progeny of type II NSCs (Fig 2A). In this system, Worniu-GAL4 drives expression in both type I and type II NSCs, while Asense-GAL80 selectively suppresses GAL4 activity in type I NSCs, thereby restricting Worniu-GAL4 activity to type II NSCs.

**Fig 2.**
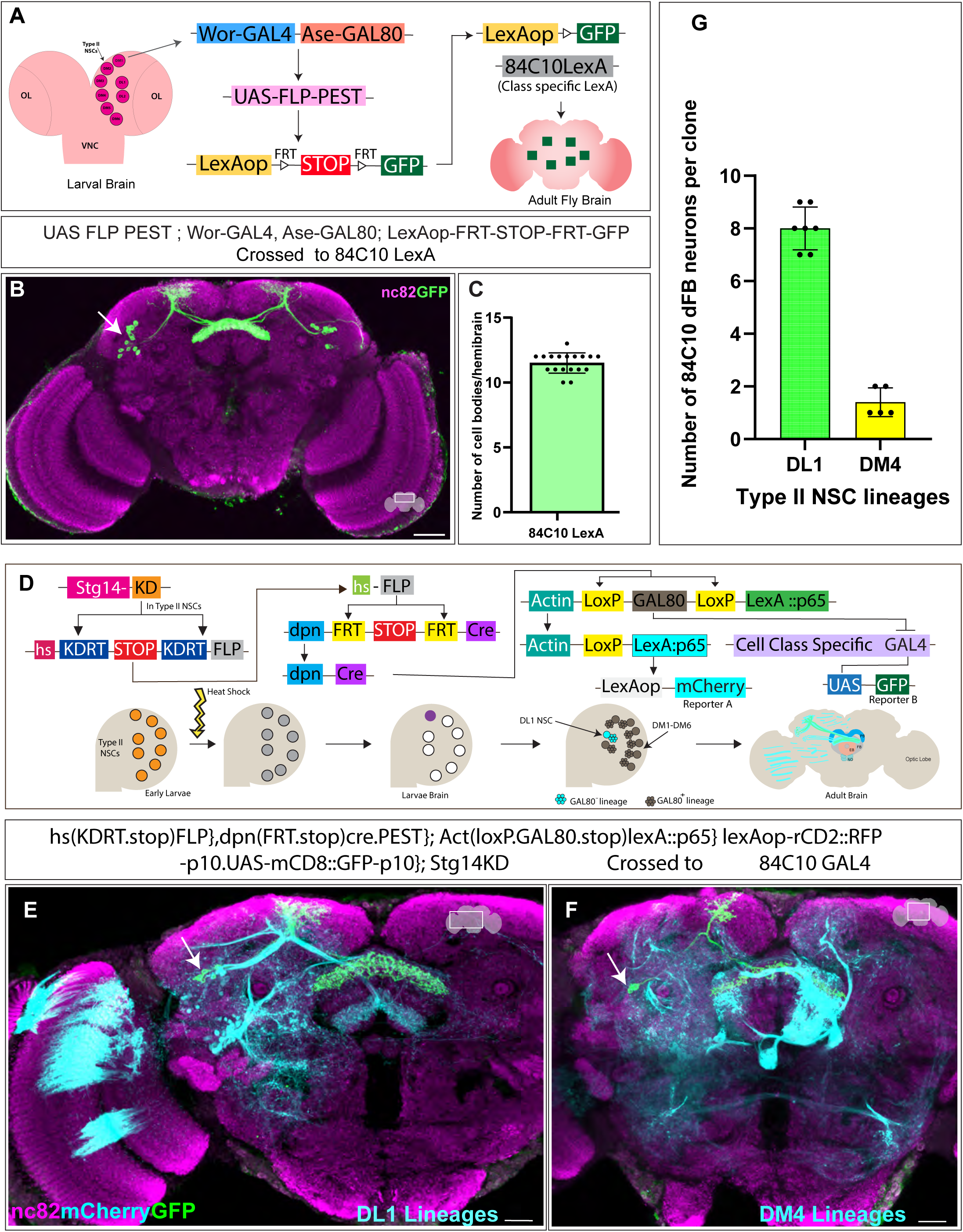
Lineage analysis reveals DL1 and DM4 Type II NSCs as sources of 84C10 dFB neurons. (A) Schematic of the Drosophila larval brain illustrating the eight Types II NSCs (magenta) per brain lobe, designated DM1–DM6 and DL1–DL2. The larval brain is subdivided into the central brain, ventral nerve cord (VNC), and optic lobes (OL). The schematic also depicts the intersectional flip-out strategy used for type II lineage analysis. In this system, Worniu-GAL4 is selectively restricted to Type II NSCs by expression of Asense-GAL80 in Type I NSCs, thereby suppressing GAL4 activity in Type I lineages. Activation of Worniu-GAL4 in Type II NSCs drives FLP recombinase expression, which excises an FRT-flanked STOP cassette, enabling expression of a LexA-dependent GFP reporter in Type II NSC derived progeny when combined with a class-specific LexA driver. (B) Representative confocal image of adult brain showing 84C10 dFB neurons derived from Type II NSCs, labeled with GFP. Axonal projections innervate the dorsal fan-shaped body (FB). The neuropil is counterstained with nc82. (C) Quantification of GFP-positive 84C10 dFB neuron cell bodies per adult hemibrain derived from Type II NSC lineages. Error bars represent mean ± SD. *n* = 18 adult hemibrains. (D) Schematic of the CLIn system (cell-class lineage intrinsic system), which combines genetic tools for lineage analysis and birthdating of neurons and glia derived from Type II NSCs. In CLIn flies, Type II NSCs express the *stg14* promoter (magenta), which drives expression of the KD recombinase. KD recombinase (orange) excises a STOP cassette flanked by KDRT sites, thereby bringing the heat-shock (*hs*) promoter into direct proximity with FLP recombinase (grey). Upon heat-shock induction, FLP is stochastically activated and excises an FRT-flanked STOP cassette, allowing recombination that brings the *dpn* promoter together with Cre recombinase (purple). Cre-mediated excision removes GAL80, resulting in the generation of an individual Type II NSC clone that is permanently positive for LexA::p65-driven *mCherry* expression (cyan) under the *actin* promoter in a lineage-specific manner. In Type II NSCs in which the heat-shock promoter is activated, *mCherry* labels all progeny derived from that NSC, including both neurons and glia. Cells that additionally express GAL4 are labeled in green in a class-specific manner, provided they are derived from the same Type II NSC lineage due to the absence of GAL80-mediated repression. Schematic adapted from Ren *et al*. (E) Composite confocal image of an adult brain showing a single DL1 Type II NSC clone induced by heat shock at 0 h after larval hatching (ALH). The DL1 clone labels the majority of 84C10 dFB neurons (green). All progeny of the DL1 Type II NSC, including both neurons and glia, are labeled in cyan (mCherry). The adult brain neuropil is counterstained with nc82 (magenta). (F) Composite confocal image of an adult brain showing a single DM4 Type II NSC clone induced by heat shock at 0 h after larval hatching (ALH). The DM4 clone labels 1-2 84C10 dFB neurons (green). All progeny of the DM4 Type II NSC, including both neurons and glia, are labeled in cyan (mCherry). The adult brain neuropil is counterstained with nc82 (magenta). (G) Quantification of the number of 84C10 dFB neurons generated per single Type II NSC clone. Each dot represents one adult brain; bars indicate mean ± SD. DL1 clones generate the majority of 84C10 neurons, whereas DM4 clones contribute a smaller subset. *n* = 5 independent clones per lineage. Scale bars, 20 μm. White arrows indicate neuronal cell bodies.

In parallel, FLP recombinase is expressed specifically in type II NSCs, where it excises an FRT-flanked STOP cassette to place LexAop-GFP in frame. As a result, GFP expression can be driven by any LexA driver exclusively in the progeny of type II NSCs. Neurons derived from type II NSCs are therefore selectively labeled when GFP is driven using a neuron class-specific LexA driver.

Using this strategy in combination with 84C10-LexA, we found that all ∼12 84C10 dFB neurons were labeled by the GFP reporter in adult brains (Figs 2B and 2C). These results demonstrate that the entire 84C10 dFB neuronal population is generated by type II NSCs.

### 84C10 dFB neurons exhibit developmental heterogeneity

Two major neuronal subclasses, columnar and tangential neurons, define the columns and layers of the FB, thereby partitioning it into distinct functional modules[42]. Previous lineage analyses have shown that among type II NSCs, the DL1 lineage is the primary contributor to FB tangential neurons, with additional minor contributions from the DM4 and DM6 lineages[24,40,43]. In contrast, columnar neurons of the CX are generated predominantly by DM1-DM4 type II NSCs[40,43]. Our previous work further demonstrated that 23E10 dFB neurons are generated mainly by DL1 NSCs, with a small contribution of 1-2 neurons from the DM1 lineage[29]. Having established that 84C10 dFB neurons also derive from type II NSCs, we next sought to define their lineage origins, which could underlie heterogeneity in their identity and function.

To identify the type II NSC lineages that generate 84C10 dFB neurons, we employed cell-class lineage analysis (CLIn), a heat shock-based lineage filtering approach that enables the generation of single type II NSC clones[44]. This method uses transient heat-inducible FLP recombinase to induce stochastic recombination events, permanently labeling the progeny of a single type II NSC with a stable mCherry reporter. Individual type II NSC lineages can be unambiguously identified in the adult brain by their characteristic morphology and connectivity patterns[24,40,43].

Heat shock was induced at 0 h after larval hatching (ALH) for 10-12 minutes at 37°C, followed by recovery and development to adulthood. Adult brains were then imaged to visualize isolated Type II NSC lineages labeled with mCherry (pseudocolored cyan) (Figs 2D, 2E, and 2F).

Using this approach, we found that the majority of 84C10 dFB neurons (∼9 cells) originated from the DL1 type II NSCs (Figs 2E and 2G). In addition, a smaller subset of 1-2 neurons was derived from the DM4 Type II NSCs (Figs 2F and 2G). These results indicate that 84C10 dFB neurons arise from two distinct type II NSC lineages rather than a single progenitor, demonstrating that this population is developmentally heterogeneous. Notably, although DL1 NSCs generate the majority of both 23E10 and 84C10 dFB neurons, distinct secondary lineages contribute to each population: DM1 to 23E10 and DM4 to 84C10.

Together, these findings establish that 84C10 dFB neurons arise from both DL1 and DM4 type II NSC lineages, revealing a lineage-based heterogeneity that may contribute to molecular and functional diversity within this neuronal population.

### Late type II NSCs give rise to 84C10 dFB neurons

We next sought to determine the birth timing of 84C10 dFB neurons. During larval development, *Drosophila* progress from 0 h after larval hatching (ALH) to ∼120 h ALH prior to pupariation, during which type II NSCs undergo sequential temporal patterning programs[5,45]. These temporal transcriptional cascades generate neuronal classes with distinct identities[9,46]. We therefore asked whether 84C10 dFB neurons are produced during early or late phases of the type II NSC temporal program.

To genetically birthdate 84C10 dFB neurons, we used the cell-class lineage analysis (CLIn) system to induce lineage labeling at defined developmental stages. we crossed 84C10-GAL4 to the CLIn background and applied heat shock at 37°C at three developmental time points: 0 h, 48 h, and 72 h ALH. Following heat shock, larvae were allowed to recover and develop to adulthood, and adult brains were subsequently examined to assess labeling of 84C10 dFB neurons corresponding to each induction window (Fig 3A).

**Fig 3.**
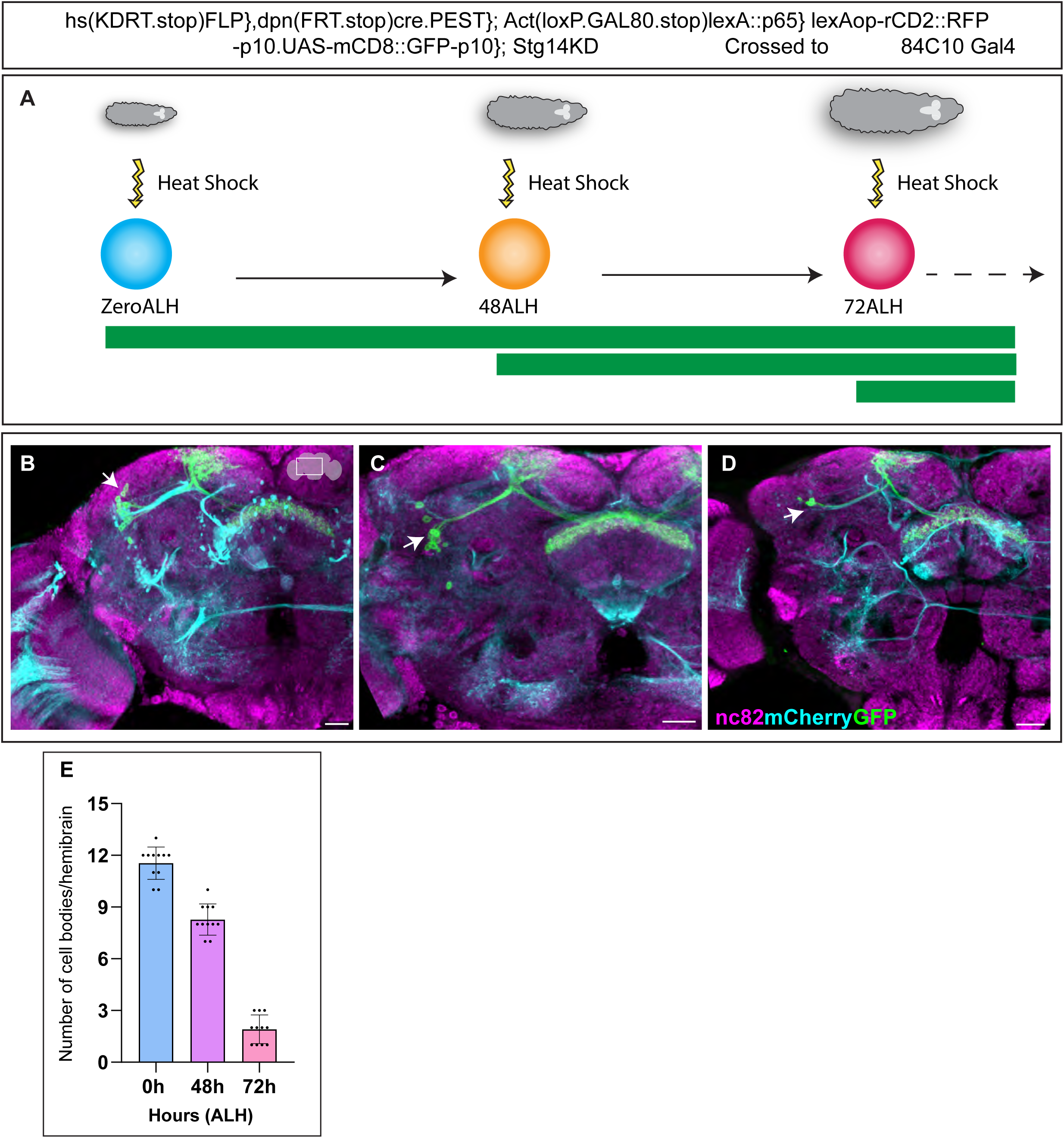
Genetic Birthdating reveals that 84C10 dFB neurons are generated during late larval development. (A) Schematic of the genetic birthdating strategy using the CLIn system to determine when 84C10 dFB neurons are generated during development. Larvae were subjected to heat shock at 37C at defined time points ALH. Yellow arrows indicate the timing of heat-shock induction, and the green bar represents the resulting window of GFP labeling. Three cohorts were heat shocked at 0, 48, or 72 h ALH, respectively. (B–D) Representative confocal images of adult brains following heat-shock induction at 0 h (B), 48 h (C), or 72 h (D) ALH, showing labeled 84C10 dFB neurons. GFP (green) marks 84C10 dFB neurons, and the adult brain neuropil is counterstained with nc82 (magenta). Heat shock was applied for 50 minutes at each developmental time point. (E) Quantification of GFP-positive 84C10 dFB neuron cell bodies per adult hemibrain following clone induction at 0, 48, and 72 h ALH. Error bars represent mean ± SD. Scale bars, 20μm. n = 16 adult hemibrains per time point. White arrows indicate neuronal cell bodies.

Heat shock induction at 0 h ALH resulted in GFP labeling of all 84C10 dFB neurons in the adult brain, confirming that this population is generated post-embryonically (Figs 3A, 3B, and 3E). When heat shock was induced at 48 h ALH, most 84C10 dFB neurons were labeled, indicating that most of this population is generated after 48 h ALH (Figs 3A, 3C, and 3E). In contrast, heat shock induction at 72 h ALH labeled only 2-3 84C10 dFB neurons in adult brains, demonstrating that a small subset of 84C10 dFB neurons continues to be generated at later stages of larval development (Figs 3A, 3D, and 3E).

Together, these results demonstrate that the majority of 84C10 dFB are generated after 48h ALH, during a developmental window that coincides with the transition from early to late temporal factors in type II NSCs (Fig 3E).

### 84C10 dFB neurons maintain expression of the late temporal transcription factor E93

Birth-dating analysis revealed that 84C10 dFB neurons are generated predominantly during the late temporal window of type II NSCs. This birth timing suggested that these neurons may inherit and maintain expression of the late temporal transcription factor E93, a key regulator of late-born neuronal identity in type II NSC lineages[5,25,29,45].

In the developing nervous system, neuronal identity is established through coordinated molecular, morphological, and functional programs, including neurotransmitter specification, connectivity, and physiological properties, and other terminal features of a neuron[15,47]. Temporal factors expressed in NSCs generate neuronal diversity by specifying distinct neuronal classes at defined developmental times[5,9,46]. Importantly, these factors can be inherited by nascent neurons and act post-mitotically to stabilize neuronal identity by regulating terminal selector genes and downstream effector pathways, including neurotransmitter and ion channel expression[14,48,49].

Given the late birth timing of 84C10 dFB neurons, we next examined whether E93 expression persists in these neurons after their birth. To assess E93 expression, we labeled 84C10 dFB neurons with GFP (Fig 4A) and stained adult brains with an antibody against E93. All 84C10 dFB neurons robustly expressed E93 in their cell bodies in the adult brain (Figs 4B, 4B′, and 4B″).

**Fig 4.**
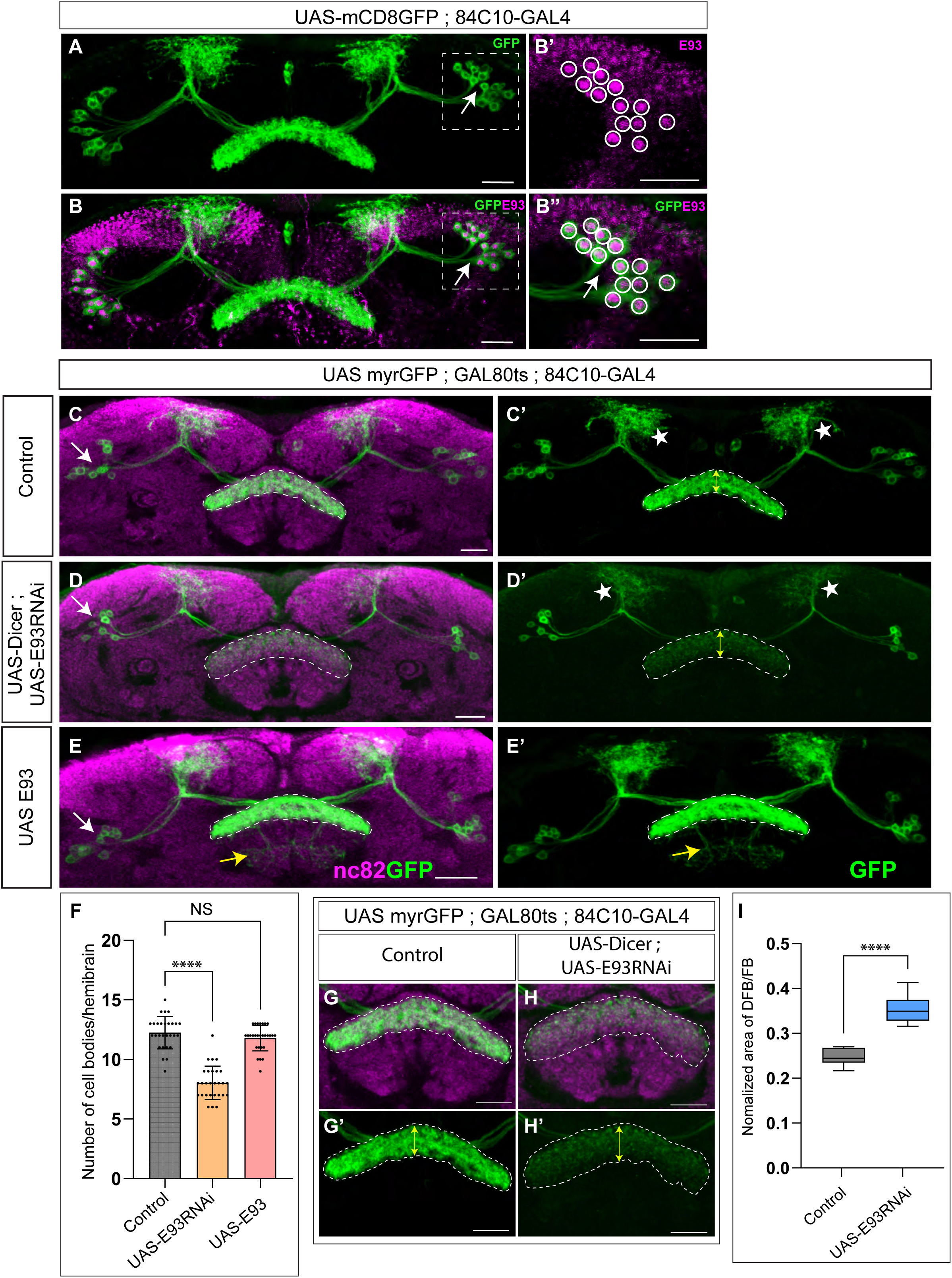
Postmitotic expression and functional requirement of the late temporal transcription factor E93 in 84C10 dFB neurons. (A) Representative confocal stack of adult brains showing 84C10 dFB neurons labeled with GFP. (B-B″) Representative confocal stacks of adult brains showing 84C10 dFB neurons (GFP, green) co-expressing the late temporal transcription factor E93 (magenta). E93 expression is maintained in the cell bodies of all 84C10 dFB neurons in adulthood. White arrows indicate neuronal cell bodies. (C-C′) Representative confocal stacks of control adult brains (KK RNAi control) showing normal number and morphology of 84C10 dFB neurons labeled with GFP. (D-D′) Representative confocal stacks of adult brains following postmitotic knockdown of E93 in 84C10 neurons using GAL80ts, showing a marked reduction in 84C10 dFB neuron number. (E-E′) Representative confocal stacks of adult brains following postmitotic overexpression of E93 in 84C10 neurons, showing largely preserved neuron number but mild axonal mistargeting. (F) Quantification of 84C10 dFB neuron cell bodies per adult hemibrain in control, E93 RNAi, and E93 overexpression conditions. Each dot represents one hemibrain; bars indicate mean ± SD. Statistical significance was assessed using one-way ANOVA followed by Dunnett’s multiple comparison test, comparing each experimental condition to the control. ∗p < 0.05, ∗∗p < 0.01, ∗∗∗p < 0.001, ∗∗∗∗p < 0.0001; NS, not significant. (G-G′) Representative confocal stacks showing layer-restricted axonal projections of 84C10 dFB neurons to fan-shaped body (FB) layers 6-7 in control adult brains. Dashed lines outline axonal projection domains. (H-H′) Representative confocal stacks showing expanded axonal innervation of 84C10 dFB neurons into ectopic middle FB layers (layers 4-5) following postmitotic E93 knockdown. (I) Quantification of normalized axonal projection area of 84C10 dFB neuronal processes relative to total FB area in control and E93 RNAi conditions. nc82 labels adult brain neuropil (magenta). Scale bars, 20μm. For expression analysis (A-B″), *n* = 10 adult hemibrains per genotype. For knockdown, overexpression, and axonal analyses (C-I), *n* = 28 adult hemibrains per genotype. White arrows indicate neuronal cell bodies.

Together, these findings demonstrate that E93 is not only inherited at the time of neuron birth but is also maintained in adult 84C10 dFB neurons. The persistent expression of E93 suggests that it may continue to regulate molecular programs required to preserve neuronal identity and functional properties throughout adulthood.

### Post-mitotic E93 knockdown causes neuronal loss and axonal expansion in 84C10 dFB neurons

Given that E93 is maintained post-mitotically in 84C10 dFB neurons (Fig 4B), we next asked whether E93 is required after neuron birth to preserve neuronal survival and identity. TTFs continue to function in postmitotic neurons, where they are thought to stabilize neuronal identity and regulate downstream effector programs. The persistent expression of E93 in adult 84C10 neurons, therefore, suggested a functional role in controlling their terminal features.

To test this, we selectively knocked down E93 in postmitotic 84C10 neurons by crossing a UAS-myrGFP ; GAL80ts ; 84C10-GAL4 to UAS-E93 RNAi. The temperature-sensitive GAL80ts repressor suppressed GAL4 activity during development, thereby restricting RNAi induction to the postmitotic phase, while myrGFP simultaneously labeled 84C10 neurons. Control animals were generated by crossing the same driver line to a KK RNAi control. Immunostaining confirmed efficient depletion of E93 protein in 84C10 positive neurons following RNAi induction (S1 Fig), demonstrating effective postmitotic knockdown within this population.

In control adult brains, all ∼12 84C10 dFB neurons were present and exhibited their characteristic morphology (Figs 4C and 4C′). In contrast, postmitotic E93 knockdown resulted in a marked reduction in the number of 84C10 neurons (Figs 4D and 4D′), quantified in (Fig 4F). In addition, we qualitatively observed that the dorsal arbor (putative dendritic region; white asterisk) appeared reduced in E93 RNAi brains (Fig D’) relative to controls (Fig C’), suggesting a possible redistribution of neurite targeting; this feature was not formally quantified. The loss of neurons suggests that E93 is required post-mitotically to maintain neuronal survival and/or neuronal fate stability.

To complement these loss-of-function experiments, we overexpressed E93 post-mitotically using the same GAL80ts-restricted system. E93 overexpression did not alter the total number of 84C10 dFB neurons but produced a mild axonal mistargeting phenotype, with some processes extending into ventral FB layers (Figs 4E, 4E’, and 4F). These findings suggest that precise E93 dosage is required for correct axonal patterning.

We next examined whether E93 depletion affected axonal targeting within the FB. In control animals, 84C10 dFB neurons restricted their projections to layers 6-7 of the FB (Figs 4G and 4G′). Interestingly, upon postmitotic E93 knockdown, axonal projections expanded markedly, extending into ectopic middle FB layers (4-5) and losing normal layer specificity (Figs 4H and 4H′). Quantification of axonal area, normalized to total FB area, revealed a significant expansion of axonal territory in E93 RNAi brains compared with controls (Fig 4I; see Methods). These data indicate that postmitotic E93 is required to maintain the layer-specific axonal lamination characteristic of 84C10 neurons. Together, these results demonstrate that postmitotic E93 is essential for maintaining the survival, identity, and precise axonal targeting of 84C10 dFB neurons.

Finally, to determine the precise timing of neuronal loss and axonal mistargeting, we examined pupal brains at 48 h and 72 h after puparium formation (APF) (S2 Fig). At 48 h APF, control animals (S2A Fig) and E93 RNAi animals (S2B Fig) showed similar numbers of 84C10 neurons, and their axonal projections were largely confined to the expected dFB layers, with no prominent axonal expansion. Quantification confirmed no significant difference in neuron number between genotypes at this stage (S1E Fig). By 72 h APF, control animals (S2C Fig) retained the normal complement of 84C10 neurons with stereotyped targeting, whereas E93 RNAi animals (S2D Fig) exhibited a clear reduction in 84C10 neuron number and a pronounced expansion of axonal innervation into ectopic

FB layers, closely resembling the adult phenotype. Quantification showed a significant decrease in neuron number in E93 RNAi animals at 72 h APF (S2E Fig).

Together, these data indicate that E93-dependent defects in neuronal maintenance and layer-specific axonal targeting emerge during late pupal neuronal maturation, rather than immediately after neuron birth or only after eclosion.

### 84C10 dFB neurons are predominantly glutamatergic but neurochemically heterogeneous

Having established the developmental origin, lineage heterogeneity, and E93 dependence of 84C10 dFB neurons, we next asked how these developmental programs are reflected in the molecular identity of mature neurons, with particular emphasis on neurotransmitter and neuropeptide expression.

To systematically assess the neurochemical identity of 84C10 dFB neurons, we employed an intersectional genetic labeling strategy. Flies carrying UAS-FLP; 84C10-LexA; LexAop-FRT-STOP-FRT-GFP were crossed to a panel of neurotransmitter- and neuropeptide-specific GAL4 driver lines. In this approach, GAL4-driven FLP excises the FRT-flanked STOP cassette, permanently activating GFP expression in 84C10 neurons that express the corresponding neurotransmitter or neuropeptide gene. Because FLP-mediated recombination is irreversible, GFP labeling reports current or prior expression of the targeted gene at any point during development or adulthood.

Using this strategy, we found that the majority of 84C10 dFB neurons were labeled by vGLUT-GAL4, indicating that these neurons are predominantly glutamatergic (Figs 5A and 5F). This finding is consistent with previous studies showing that many dorsal fan-shaped body neurons use glutamate as a primary fast-acting transmitter[22,33,50]. Recent work has shown that 23E10 dFB neurons are neurochemically heterogeneous, including small non-glutamatergic subsets[36].

**Fig 5.**
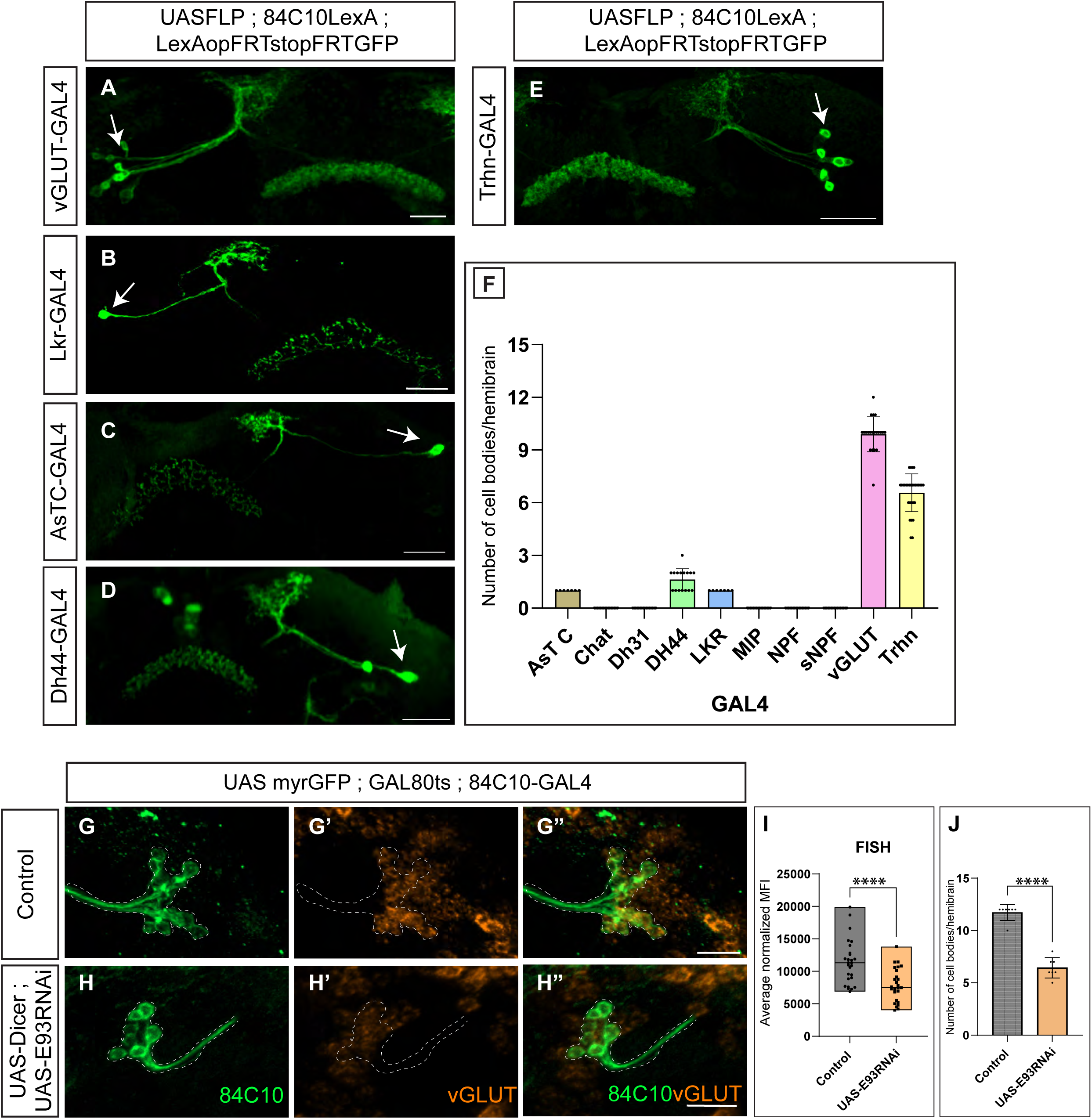
84C10 dFB Neurons Are Predominantly Glutamatergic and Require Post-mitotic E93 to Maintain Neurotransmitter Identity. (A-E) Intersectional genetic labeling strategy to assess neurotransmitter and neuropeptide expression in 84C10 dFB neurons. Flies carrying *UAS-FLP; 84C10-LexA; LexAop-FRT-STOP-FRT-GFP* were crossed to neurotransmitter- or neuropeptide-specific GAL4 driver lines. GAL4-driven FLP excision permanently activates GFP expression in 84C10 neurons that express the corresponding marker at any point during development or adulthood. (A) vGLUT-GAL4 labels the majority of 84C10 dFB neurons, indicating a predominantly glutamatergic identity. (B-D) Smaller subsets of 84C10 dFB neurons are labeled by Dh44-GAL4 (B), AstC-GAL4 (C), and Lkr-GAL4 (D), revealing neuropeptidergic heterogeneity within the population. (E) Trh-GAL4 labels a substantial subset of 84C10 dFB neurons, indicating the capacity for serotonergic biosynthesis. (F) Quantification summarizing the number of 84C10 dFB neurons labeled by each neurotransmitter or neuropeptide driver. Most neurons are vGLUT-positive, with smaller subsets expressing Dh44, AstC, Lkr, or Trh. Drivers for Dh31, MIP, NPF, sNPF, and ChAT did not label 84C10 neurons. n = 10 adult hemibrains per genotype. (G-G″) Representative confocal images showing control adult brains in which 84C10 dFB neurons are labeled with myrGFP (green) and analyzed by fluorescent in situ hybridization (FISH) for vGLUT mRNA. Robust vGLUT expression is detected in neuronal cell bodies. (H-H″) Representative confocal images of adult brains following postmitotic E93 knockdown in 84C10 dFB neurons (*UAS-myrGFP; GAL80ts; 84C10-GAL4 > E93 RNAi*). vGLUT mRNA signal is markedly reduced in surviving 84C10 neurons compared with controls. (I) Quantification of vGLUT fluorescence intensity in individual 84C10 dFB neuron cell bodies. Postmitotic E93 knockdown significantly reduces vGLUT expression relative to controls, indicating loss of glutamatergic identity in surviving neurons. (J) Quantification of total 84C10 dFB neuron number per adult hemibrain in the same animals analyzed in (G-I). Postmitotic E93 knockdown results in a significant reduction in neuron number. Error bars represent mean ± SD. For neuronal analyses (I, J), n = 14 adult hemibrains per genotype. Statistical significance was assessed using unpaired two-tailed Student’s t-tests with Welch’s correction. Asterisks denote significance levels as indicated. Scale bars, 20 μm. White arrows indicate neuronal cell bodies.

In addition, smaller subsets of 84C10 neurons were labeled by Dh44-GAL4 (approximately three cells), AstC-GAL4 (one cell), and Lkr-GAL4 (one cell), revealing neuropeptidergic diversity within the 84C10 dFB population (Figs 5B-5D and 5F).

Notably, we also observed that approximately seven 84C10 dFB neurons were labeled by Trh-GAL4 (Figs 5E and 5F), which drives expression of tryptophan hydroxylase (Trh), the rate-limiting enzyme required for serotonin (5-HT) biosynthesis. Previous studies have implicated serotonergic signaling in modulating dFB function primarily through receptor expression in this region[51,52]. However, serotonergic biosynthetic capacity within genetically defined dFB neuron populations has not been previously reported. Our data, therefore, indicate that a substantial subset of 84C10 neurons can engage the serotonergic biosynthetic pathway at some point during development or adulthood.

In contrast, GAL4 drivers for Dh31, MIP, NPF, sNPF, and ChAT did not label 84C10 dFB neurons, indicating that these neuropeptides and the cholinergic neurotransmitter pathway are not characteristic features of this population (Fig 5F).

Together, these results demonstrate that 84C10 dFB neurons exhibit a restricted yet heterogeneous neurochemical profile, characterized by a predominantly glutamatergic identity with additional diversity contributed by serotonergic and select neuropeptidergic features. This molecular heterogeneity parallels the developmental heterogeneity revealed by lineage tracing and birth-dating experiments and provides a framework for understanding how E93-dependent temporal programs may shape the functional properties of this neuronal population.

After defining the developmental origin, lineage heterogeneity, and E93 dependence of 84C10 dFB neurons, we next asked whether their neurotransmitter identity is maintained in adulthood. Because the FLP-based intersectional strategy used above captures expression history rather than exclusively postmitotic expression, we directly examined neurotransmitter expression in mature 84C10 neurons.

To assess postmitotic neurotransmitter identity, we performed fluorescent in situ hybridization (FISH) on adult brains using probes against vGLUT, the vesicular glutamate transporter. FISH analysis revealed robust vGLUT mRNA expression in 84C10 dFB neurons in the adult brain (S3A-S3A” Figs), demonstrating that these neurons maintain a glutamatergic identity post-mitotically.

We next examined whether neuropeptidergic markers identified using genetic reporters are maintained post-mitotically in the neuronal population shared between the 84C10 and 23E10 driver lines. This shared population represents the core subset of dFB neurons common to both drivers. Using FISH on adult brains, we assessed expression of vGLUT, AstC, and Dh44 mRNAs in the shared 84C10-23E10 neurons. Consistent with our main findings, shared neurons robustly expressed vGLUT mRNA, confirming their glutamatergic identity in adulthood (S4A-S4D Figs).

In contrast, Dh44 and AstC mRNA were not detected in the shared neuron population under the conditions tested (S4A, S4B, S4D Figs and S4E, S4E’, S4E’’ Figs). These results suggest that glutamatergic identity is a stable postmitotic feature of the shared dFB neurons, whereas AstC and Dh44 expression may be restricted to non-overlapping subsets of 84C10 neurons, to transient developmental stages, or to expression levels below the detection threshold of this assay.

### E93 regulates vGLUT expression in 84C10 dFB neurons

Given that glutamatergic identity is a stable postmitotic feature of 84C10 dFB neurons, we next asked whether E93 is required to maintain this neurotransmitter identity after neuronal birth. Our earlier analyses showed that postmitotic knockdown of E93 reduces the number of 84C10 neurons and disrupts axonal targeting. We therefore examined whether loss of E93 also affects expression of the glutamatergic effector gene *vGLUT* in the surviving neurons.

To test this, we selectively knocked down E93 post-mitotically in 84C10 neurons using genotype UAS-myrGFP ; GAL80ts ; 84C10-GAL4, which temporally restrict RNAi induction to postmitotic stages. Flies were reared at the permissive temperature during development and shifted to the restrictive temperature in adulthood to induce E93 RNAi, while myrGFP simultaneously labeled 84C10 neurons.

To directly assess neurotransmitter identity, adult brains were analyzed by FISH using probes targeting *vGLUT* mRNA, enabling single-cell-resolution measurement of glutamatergic gene expression. In control animals, 84C10 dFB neurons exhibited robust *vGLUT* mRNA expression in their cell bodies (Figs 5G-5G’’ and 5I). In contrast, postmitotic E93 knockdown resulted in a marked reduction of *vGLUT* mRNA levels in the 84C10 neurons (Figs 5H-5H’’ and 5I).

To quantify this effect, we measured *vGLUT* fluorescence intensity in individual 84C10 neuron cell bodies and averaged values across multiple brains. This analysis revealed a significant decrease in *vGLUT* signal intensity in E93 RNAi animals compared with controls (Fig 5I), indicating that E93 is required to maintain glutamatergic identity in postmitotic 84C10 neurons.

Consistent with our earlier findings, postmitotic E93 knockdown also resulted in a significant reduction in the total number of 84C10 dFB neurons in the same animals (Fig 5J). Importantly, reduced vGLUT expression was observed within surviving neurons, indicating that loss of glutamatergic identity is not solely a secondary consequence of neuron loss.

Together, these results demonstrate that E93 acts post-mitotically to sustain expression of the glutamatergic effector gene *vGLUT* in 84C10 dFB neurons, linking late temporal transcriptional programs to the maintenance of neurotransmitter identity in the 84C10 dFB neuron population.

### E93 is required in 84C10 dFB neurons for metabolic adaptation

The dFB neurons have been broadly implicated in the regulation of behavioral state and metabolic homeostasis[31,32,53]. Because we found that postmitotic loss of E93 disrupts the survival, wiring, and neurotransmitter identity of 84C10 dFB neurons, we asked whether impairing E93-dependent maintenance of this dFB population alters organismal metabolic responses under dietary challenge.

To address this question, we quantified triglyceride (TAG) accumulation in adult flies following exposure to a well-established diet-induced obesity model[54]. Control flies and flies with postmitotic E93 knockdown in 84C10 dFB neurons were transferred 4-5 days after eclosion to either a control diet (CD) or a 30% high-sugar diet (SD) and maintained on the respective diets for seven days, with food replaced every other day. TAG levels were then measured using a colorimetric assay and normalized to protein content, as previously described[55].

Under control diet conditions, TAG levels were comparable between control and E93 RNAi animals, indicating that basal lipid storage was not substantially affected by postmitotic loss of E93 (S5 Fig). In contrast, following exposure to a high-sugar diet, control flies exhibited a robust increase in TAG accumulation, consistent with diet-induced lipid storage. Strikingly, flies with postmitotic E93 knockdown in 84C10 dFB neurons failed to accumulate TAGs, displaying significantly lower TAG levels than controls on the same diet (S5 Fig).

These results indicate that postmitotic E93 activity in 84C10 dFB neurons is required for fat accumulation in response to a sugar-rich diet. Although we did not directly measure food intake in these animals, prior studies have shown that fan-shaped body neurons regulate state-dependent nutrient handling rather than feeding capacity per se. In this context, our finding that loss of E93 disrupts neuron number, axonal targeting, and glutamatergic identity in this population is consistent with both the reported roles of dFB neurons and the metabolic phenotypes we observe here. Together, these data suggest that E93-dependent maintenance of 84C10 dFB neuron identity is required not only for proper circuit architecture, but also for translating dietary sugar into appropriate energy-storage responses.

## Discussion

Our work defines the developmental origins and developmental heterogeneity of the dFB neuron subtypes. Using lineage-restricted labeling and CLIn-based clonal analysis, we show that the full 84C10 dFB population is generated from type II NSCs, with most neurons arising from the DL1 lineage and a smaller contribution from DM4, and that they are produced primarily during a late larval window (after ∼48 h ALH). These findings are consistent with the prolonged temporal patterning of type II NSCs and their contribution to central-complex diversity[22,24,34].

The partial but incomplete overlap between 84C10 and 23E10 labeling supports the idea that “dFB neurons” comprise multiple related subtypes rather than a single uniform class. This interpretation is supported by our clonal analysis and by recent connectomic and transcriptomic analyses, which show that CX circuits consist of closely intermingled neuron types that share gross projection patterns but differ in lineage origin, molecular identity, and likely function.

Functionally, such heterogeneity may be well suited for state-dependent behaviors. dFB-related circuits have been implicated in selecting food options under sensory conflict, in which hunger and internal-state signals converge onto fan-shaped body neurons to guide adaptive choice[31,32]. It is possible that lineage-specific subsets of the dFB neurons, such as 84C10-labelled neurons in DM4 and 23E10-labelled neurons in DM1, engage in ubiquitous behaviors. Previous work has shown a pair of neurons regulating internal state and sleep behavior[32,52,57–59]. Whether the lineage identity reflects functional diversity requires testing.

A central finding of our study is that the late TTF, E93, is not merely a developmental fate specifier; it is expressed in adult 84C10 neurons and governs their terminal identity. Post-mitotic knockdown of E93 disrupts layer-restricted axonal targeting, leading to expansion into ectopic FB layers, and reduced expression of the glutamatergic effector gene vGLUT. Importantly, these defects emerge during late pupal maturation (∼72 h APF), rather than immediately after neuronal birth, indicating a role for E93 in late-stage neuronal maturation, maintenance, and circuit refinement.

A plausible explanation for the ectopic delamination phenotype is the disruption of repulsive cues that normally confine dFB neurons to layers 6-7. Semaphorin-mediated repulsion has been shown to ensure precise laminar targeting of these neurons[60], suggesting that E93 functions upstream of this pathway. In this model, E93 may coordinate terminal differentiation with spatial positioning by regulating the expression of semaphorin ligands or receptors, thereby coupling temporal identity to laminar specificity. Alternatively, E93 may act more broadly to stabilize neuronal adhesion or cytoskeletal programs required for layer-restricted targeting. Dissecting whether E93 directly interfaces with semaphorin signaling or instead governs lamination through parallel developmental mechanisms will be essential for understanding how temporal transcription factors translate lineage history into precise circuit wiring.

These findings place E93 within a growing class of TFs that bridge development and adulthood by stabilizing terminal neuronal identity programs, including neurotransmitter specification and connectivity[13,14,48]. Although E93 has not traditionally been classified as a terminal selector, our data suggest that it performs terminal selector-like functions in mature dFB neurons. Previous studies on E93 have primarily focused on NSCs, in which it has been shown to play essential roles in fate specification and proliferation[29,61].

We find that 84C10 dFB neurons are predominantly glutamatergic, with robust adult vGLUT expression confirmed by FISH. Reporter-based analyses suggest additional molecular heterogeneity within this population, including small subsets expressing neuropeptides such as Dh44, AstC, and Lkr, as well as serotonergic biosynthetic capacity via Trh. A close interpretation is that glutamate mediates fast output from this dFB module, whereas peptidergic and neuromodulatory components tune gain and state dependence, as proposed for other fan-shaped body circuits[31,33]. The requirement for E93 to maintain vGLUT expression directly links a TTF to an adult neuronal function. Disruption of neurotransmitter identity together with loss of layer-specific targeting provides a mechanistic explanation for how circuit output is impaired following postmitotic E93 loss. Given prior work showing that FB neurons encode food choice by integrating hunger state and sensory information, such disruptions are expected to impair the translation of internal state into adaptive feeding decisions[31,32]. The mechanism by which E93 regulates vGLUT expression warrants further study.

Postmitotic E93 knockdown in 84C10 dFB neurons selectively disrupts metabolic adaptation: flies fail to appropriately accumulate triglycerides on a high-sugar diet, while baseline lipid levels on a control diet remain largely intact. This phenotype supports a model in which maintaining identity and wiring within a small, genetically defined dFB population is required for state-dependent metabolic responses under dietary challenge, rather than for basal metabolic homeostasis. This finding aligns with broader roles for E93 in adult physiology. Pan-neuronal E93 knockdown has been shown to alter feeding-related behaviors, circadian rhythms, and systemic metabolic state, consistent with E93 functioning as a hormone-responsive transcriptional regulator that supports adult behavioral and metabolic programs[56].

Our findings resonate with work on mammalian orthologs of E93, ligand-dependent nuclear receptor corepressor-like (LCORL). In mice, loss of LCORL results in reduced body size, altered growth trajectories, and broad metabolic changes, including protection against diet-induced obesity and improved glucose homeostasis[62].

Together, these studies suggest that the E93/LCOR/LCORL axis represents an evolutionarily conserved transcriptional module linking hormonal cues, neuronal function, and long-term metabolic physiology across species.

### Model and future directions

We propose a model in which late-born 84C10 dFB neurons inherit and maintain E93, and continued E93 activity stabilizes neuronal survival, layer-specific wiring within the fan-shaped body, and the glutamatergic effector program. Loss of E93 post-mitotically leads to erosion of neuronal identity, circuit disorganization, and failure to support normal metabolic adaptation under dietary challenge. Uncovering the mechanisms by which E93 regulates neuronal terminal features represents an important avenue for future investigation.

Importantly, while postmitotic E93 knockdown disrupts diet-induced triglyceride accumulation, we cannot yet distinguish whether this reflects altered feeding behavior, impaired internal-state processing, or a direct metabolic defect. These possibilities are not mutually exclusive and will require feeding-rate and food-choice assays, as well as metabolic flux measurements, in future work.

## Materials and methods

### *Drosophila* stocks and rearing

*Drosophila melanogaster* adult flies (both male and female flies) and larvae were maintained and raised on standard Bloomington Food formulation at 25°C in a humidity-controlled incubator under a 12:12h LD cycle. The RNAi and FLPStop experimental flies were raised at 29°C.

For all feeding experiments, flies were collected under CO2 anesthesia 2–4 days following eclosion and housed in groups of 20-40 within culture vials to age until 4-5 days old. As a control, we used *w1118 Canton-S (wCS)* flies (gift from Anne Simon, University of Western Ontario), which were obtained by backcrossing a *w1118* strain (Benzer lab, Caltech) to *Canton-S (CS)* (Benzer lab, Caltech) for 10 generations.

Parental flies used in this work are listed in the S1 Table along with RRIDs.

### Immunostaining

5-7 day old adult flies were decapitated, and brains were dissected in cold Schneiders insect media (Sigma Aldrich) and then fixed at room temperature for 27 minutes in 4% paraformaldehyde (PFA) (EMS) in PBST (1X phosphate-buffered saline with 0.5 % Triton X-100). After fixation, three quick washes were given to brains followed by three 20-minute washes in 1X PBST (the long washes or antibody staining was done over the nutator). The brains were incubated for 40 minutes at room temperature in a block solution (1X PBS with 0.5% Triton X-100 with the addition of 2.5% Normal Goat serum and 2.5% Normal Donkey Serum) (Jackson ImmunoResearch). Following blocking, brains were incubated for 2 nights at 4°C with primary antibodies (primary antibodies are made in block solution). Subsequently, the brains were given three quick washes and three long washes for 20 minutes in PBST. The Brains were further incubated with secondary antibody for two nights at 4°C. After the secondary antibody, the brains were given three quick washes again, and three 20-minute PBST washes.

After antibody staining, the brain samples were subsequently mounted. The brain samples were washed with PBS for 10 minutes to remove the detergent (Triton). The detergent interferes with brain samples sticking to poly-lysine-coated cover slips. The brain samples were mounted on a poly-lysine-coated cover slip and then dipped in MilliQ water for 2 seconds to remove salts. Following this, brain samples were dehydrated in jars containing increasing concentrations of alcohol: 30%, 50%, 75%, 95%, 100%, 100%, and 100% (each step lasted 10 minutes). The alcohol dehydration was followed by clearing the brain samples in three successive jars of xylene (each step lasts 5 minutes). The brain samples were finally mounted in a DPX mounting medium (Sigma-Aldrich #06522). The slides were allowed to dry for three days before imaging. Janelia’s protocol provides additional information on DPX mounting.

The following primary antibodies were used: chicken anti-GFP (1:1500), mouse anti-nc82 (1:50), guinea pig anti-E93 (1:300), and rabbit anti-RFP (1:500). Fluorescent in situ hybridization probes (Molecular Instruments) were used to detect *vGlut*, *DH44*, and *AstC* transcripts.

**Pupal brains:** For pupal brain analysis, 0 h pupae were marked and dissected at 48 h and 72 h APF in 1× PBS, following a previously described pupal brain dissection protocol[63]. Pupal brains were fixed overnight at 4°C in acid-free glyoxal containing 5% sucrose. All subsequent blocking, immunostaining, dehydration, and mounting steps were performed as described above for adult brains.

### RNA in situ hybridization

Adult brains were dissected in Schneider’s S2 medium and transferred to 500 µL tubes.

Brain samples were washed once in PBS for 1 min and then fixed in 4% paraformaldehyde containing 0.5% Triton X-100 in PBS for 25 min. Brains were subsequently washed twice in RNase-free 0.5% PBST for 20 min each with gentle nutation. Brains were pre-equilibrated in 300 µL of pre-heated HCR HiFi probe hybridization buffer at 37°C for 1 h, followed by incubation with HCR probes (Molecular Instruments) at a final concentration of 5-10 nM overnight at 37°C with nutation. After hybridization, brains were washed four times in pre-heated HCR HiFi probe wash buffer at 37°C for 20 min each. Samples were then incubated in 300 µL amplification buffer at room temperature for 30 min. Imager hairpins (attached with Dyes) were individually heated to 95°C for 90 s and immediately snap-cooled to 4°C. Five microliters of each hairpin were added to the brains in amplification buffer, and the mixture was incubated overnight at room temperature with nutation.

Finally, brains were washed twice in 2× SSC buffer for 30 min each, followed by a wash in 2× SSC for at least 10 min at room temperature. Samples were mounted on poly-L-lysine-coated coverslips in ProLong Diamond mounting medium (Thermo Fisher Scientific) for imaging.

### Microscopy and image analysis

Fixed Fluorescent image stacks of fly brains were taken on a LSM 780 and 980 confocal microscope (Zeiss) using a 40x water immersion objective. Optical sections were acquired at 1.0 μm and most images were taken at 0.6X zoom in a 1024X1024 configuration. In the figures, only regions of interest from fixed images were shown like slices corresponding to the FB neuropil (stained with nc82) and the slices that give the full projection of 84C10 dFB neurons stained with GFP. Images were analyzed and processed using Fiji (ImageJ), and Adobe Photoshop. The final figures were composed using Adobe Illustrator.

### Clonal Analysis and Birth dating

The virgin females from the Clin stock were mated with males from the neuron class-specific GAL4 line. Egg laying was performed on agar-apple juice plates. The eggs were collected for a 0-3 h window, and the process was repeated multiple times. The 0-3 h-old eggs were allowed to hatch on regular food. For clonal analysis (lineage analysis), 0-3 h-old, hatched larvae were heat shocked at 37°C for 10-12 minutes, then left on regular food to develop into adults. For the birth-dating experiment, 0-3 h-old larvae were heat-shocked for 50 minutes at 0 h, 48 h, and 72 h. The larvae were allowed to grow on regular food till adult. For both lineage analysis and birth dating, the adult flies aged 5-7 days were dissected.

### Dietary manipulations

Flies were transferred to vials containing respective diets 4-5 days after eclosion and left on their food for 7 days; fresh food was provided every other day. The composition and caloric amount of each diet were as follows:

-”Control Diet/CD” was a standard cornmeal food (Bloomington Food B recipe), with approx. 0.6 cal/g.

-”Sugar Diet/SD” was 30 g of table sugar added to 89 g Control Diet for 100 mL final volume of 30% sucrose w/v, with approx. 1.4 cal/g.

### Triacylglyceride (TAGs) Assay

The levels of TAG and protein were measured as previously described in Tennessen et al, 2014; n=1 equals 2 flies.

### Quantification and Statistical Analysis

Neuron cell body counting was done using a cell counter in ImageJ. The statistical analysis and graphs were made in GraphPad Prism. Student’s t-test or one-way ANOVA provided P values. The one-way ANOVA followed Dunnett’s Test, or Tukey’s Test, or otherwise stated. Error bars represent as mean ±SD and asterisks indicate levels of significance (*: p<0.05, **: p<0.01, ***: p<0.001, ****: p<0.0001, NS, non-significant).

n represents the sample size.

To quantify the 84C10 neurons’ axonal expansion upon post-mitotic knockdown of E93 (Figure 4), we performed area analysis using ImageJ. We analyzed a single optical section of adult brains, measuring the total area of 84C10 dFB neuropil and normalizing it to its respective FB area. This was done for both control and experimental brain samples. The normalized areas were statistically compared between control and experimental groups using a t-test in GraphPad Prism.

Antibodies, software and Chemical information used in this work can be found in S2 Table

## Acknowledgments

We thank Chris Doe and Durafshan Sakeena Syed for reading the manuscript and providing feedback. Claude Desplan, Tzumin Lee, and Gerry Rubin for providing the reagents. The research was supported by the Sloan Research Fellowship FG-2023-20617, 2023 BBRF Young Investigator Grant, McKnight Scholars Award, National Science Foundation CAREER Award IOS-2047020, Air Force Office of Scientific Research FA9550-24-1-0214, and NINDS R01NS136555 to MHS; and by NIH R01DK130875, the Klingenstein-Simons Fellowship in the Neurosciences, the Rita Allen Foundation (all to M.D.). Stocks obtained from the Bloomington *Drosophila* Stock Center (NIH P40OD018537) were used in this study. The monoclonal nc82 antibody was obtained from the Developmental Studies <u>Hybridoma</u> Bank, created by the NICHD of the NIH and maintained at the University of Iowa, Department of Biology, Iowa City, IA 52242.

## Author Contributions

Conceptualization: A.R.W and M.H.S.; methodology: A.R.W.; visualization: A.R.W. and M.H.S.; writing-original draft: A.R.W.; editing: M.H.S.; funding acquisition: M.H.S. and supervision: M.H.S.; FISH Experiments R.A.; Pupae Experiments by Z.M, Q.B.J, and T.A.; Metabolic experiments by R.W and M.D.; Few Neurochemical identity experiments by M.V.

---

**S1 Fig.**
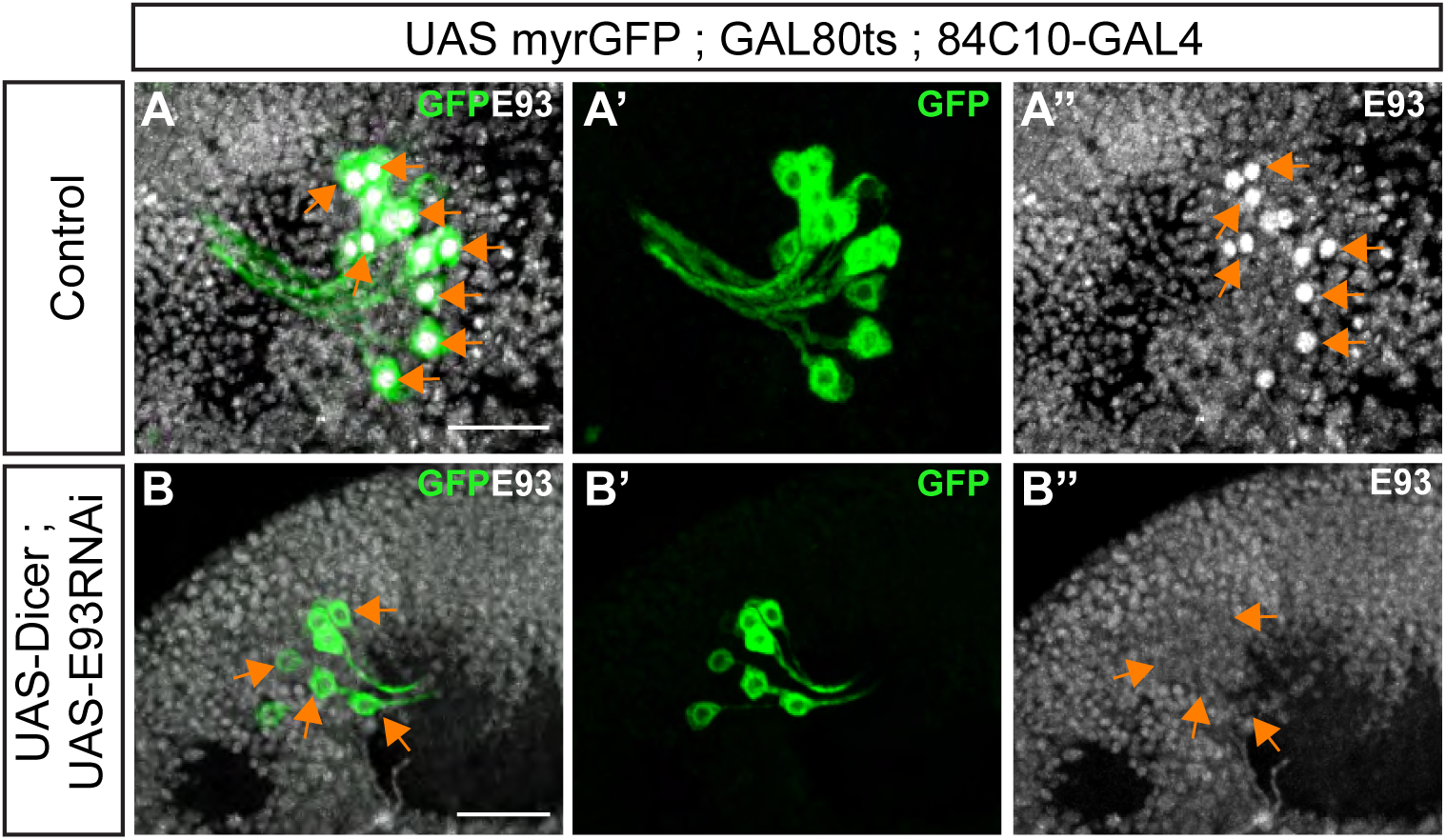
84C10-GAL4 mediated E93 RNAi efficiently depletes E93 in postmitotic 84C10 neurons. (A-A″) In control adult brains, 84C10 positive neurons labeled with GFP (green) co-express E93 protein (gray). (A) Merged image showing co-localization of 84C10-GFP and E93. (A′) 84C10-GFP (A″) E93 immunostaining. (B-B″) Postmitotic knockdown of E93 in 84C10 neurons using 84C10-GAL4 > UAS-E93 RNAi. (B) Merged image showing 84C10-GFP labeled neurons (green) with loss of E93 signal (gray). (B′) 84C10-GFP. (B″) E93 immunostaining. Orange arrows indicate 84C10-positive cell bodies. E93 signal is detected in 84C10 neurons in controls but is absent following RNAi expression. Scale bar, 20 μm. n = 10 brains.

**S2 Fig.**
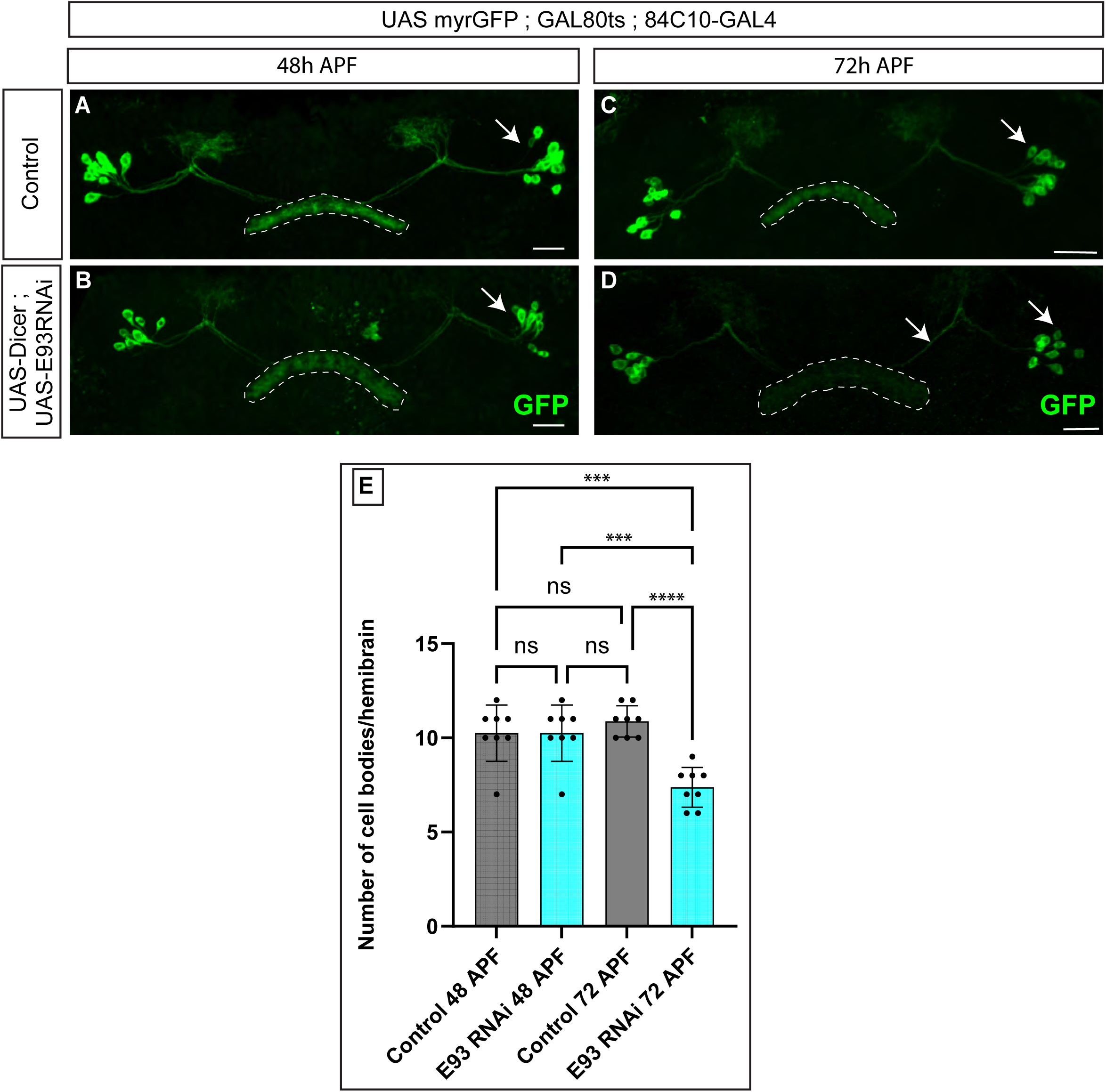
E93-dependent neuronal loss and axonal mistargeting emerge during late pupal maturation. (A, B) Representative confocal images of 84C10 dFB neurons in control (A) and postmitotic E93 RNAi (B) pupal brains at 48 h after puparium formation (APF). At this stage, the number of 84C10 neurons and their axonal targeting within the fan-shaped body (FB) are indistinguishable between control and E93 RNAi animals. (C, D) Representative confocal images of 84C10 dFB neurons in control (C) and postmitotic E93 RNAi (D) pupal brains at 72 h APF. At this later stage, E93 knockdown animals exhibit a pronounced reduction in 84C10 neuron number accompanied by aberrant axonal projections and expansion into inappropriate FB layers, closely resembling the adult phenotype. (E) Quantification of 84C10 dFB neuron number per hemibrain at 48 h and 72 h APF in control and E93 RNAi animals. Neuron numbers are comparable between genotypes at 48 h APF but are significantly reduced in E93 RNAi animals at 72 h APF. Statistical analysis was performed using one-way ANOVA followed by Tukey’s multiple comparisons test. n= 6 adult brains. Error bars represent mean ± SD. Scale bars, 20μm. ∗p < 0.05, ∗∗p < 0.01, ∗∗∗p < 0.001, ∗∗∗∗p < 0.0001; NS, not significant. Dashed outlines indicate the area of projection from 84C10 neurons.

**S3 Fig.**
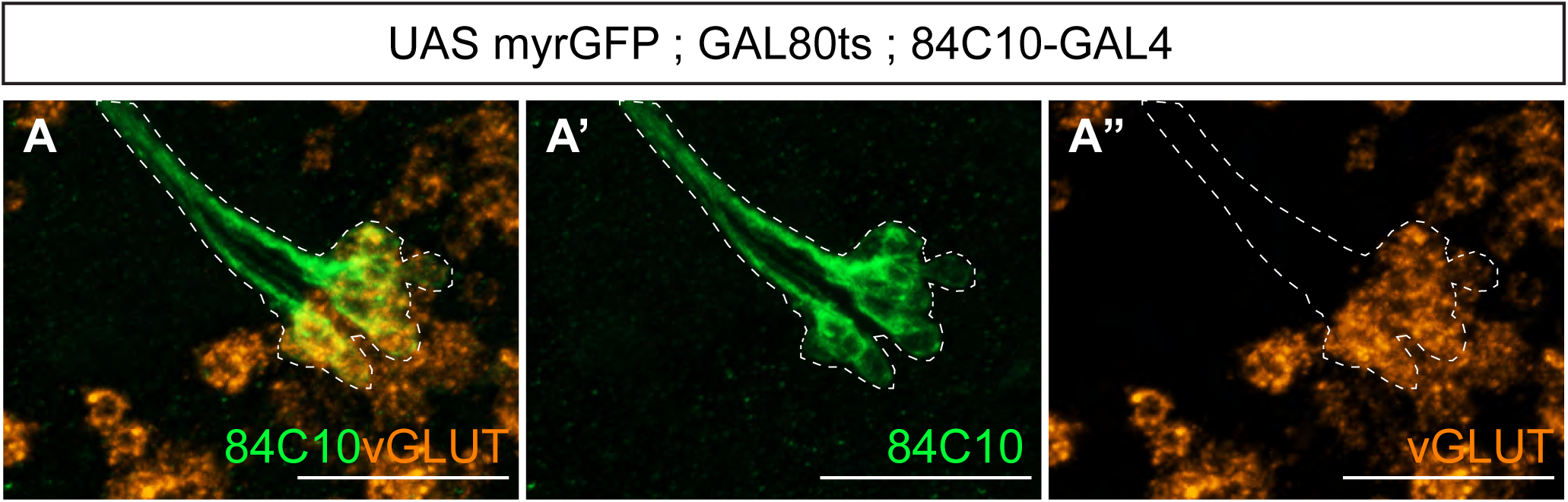
vGLUT FISH Validates Glutamatergic Identity of 84C10 dFB Neurons. (A-A″) Fluorescent in situ hybridization (FISH) for vGLUT mRNA in adult brains with 84C10 dFB neurons labeled by GFP reveals robust postmitotic vGLUT expression in neuronal cell bodies, confirming maintenance of glutamatergic identity in adulthood. Scale bars, 20 μm. n = 6 adult hemibrains per genotype. Dashed outlines indicate 84C10 dFB neuron cell bodies.

**S4 Fig.**
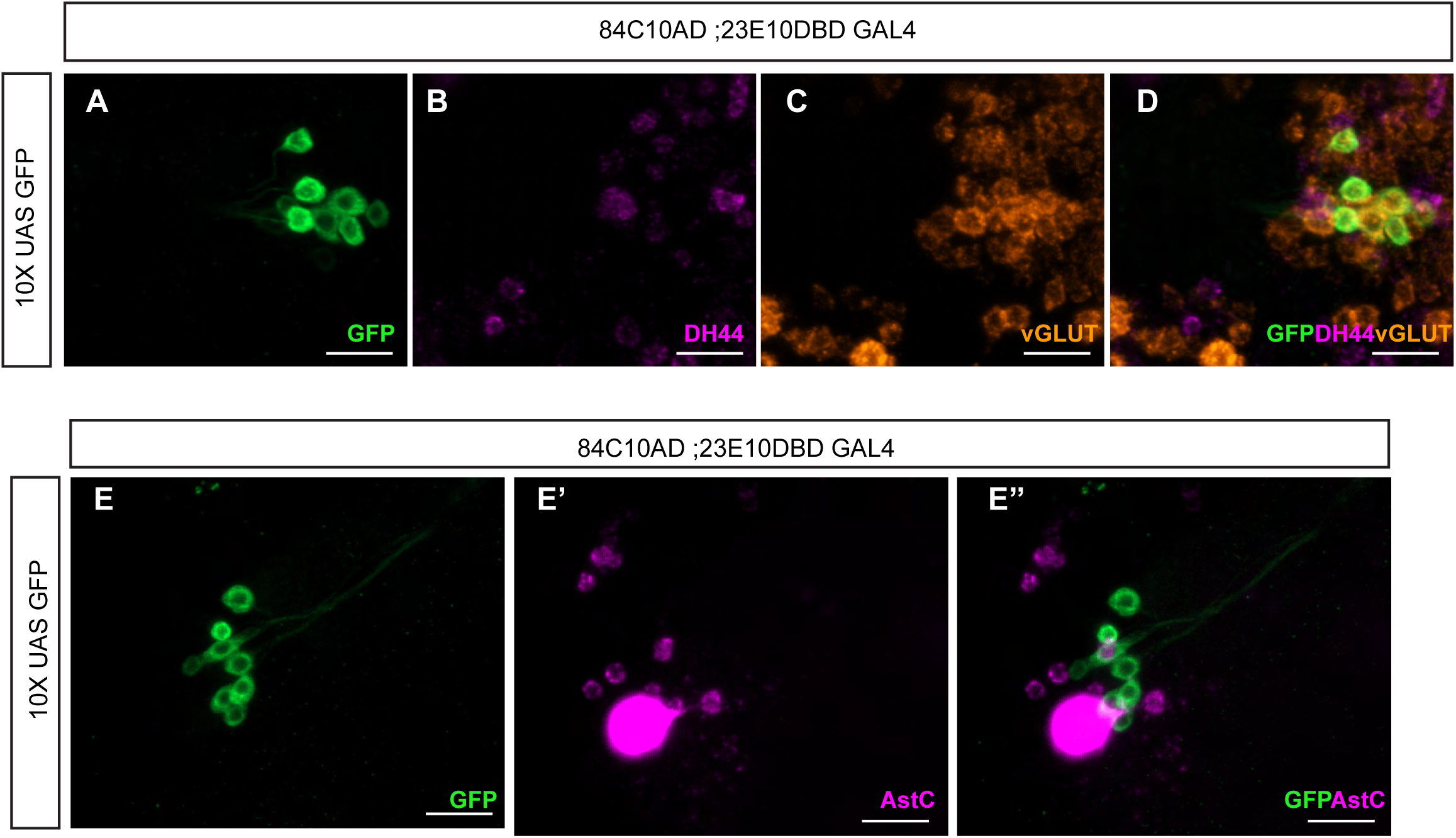
Shared 84C10-23E10 dFB neurons maintain glutamatergic identity in adulthood. (A-D) Representative confocal images showing shared 84C10-23E10 neurons labeled with GFP (A), probed for Dh44 mRNA (B), vGLUT mRNA (C), and the merged image (D). vGLUT mRNA is robustly detected in GFP-positive shared neurons, confirming their glutamatergic identity in the adult brain, whereas Dh44 mRNA is not detected in this population. (E-E″) Representative confocal images showing shared 84C10-23E10 neurons labeled with GFP (E), probed for AstC mRNA (E′), and the merged image (E″). AstC mRNA is not detected in GFP-positive shared neurons under the conditions tested. Scale bars, 20 μm. n= 6 hemibrains.

**S5 Fig.**
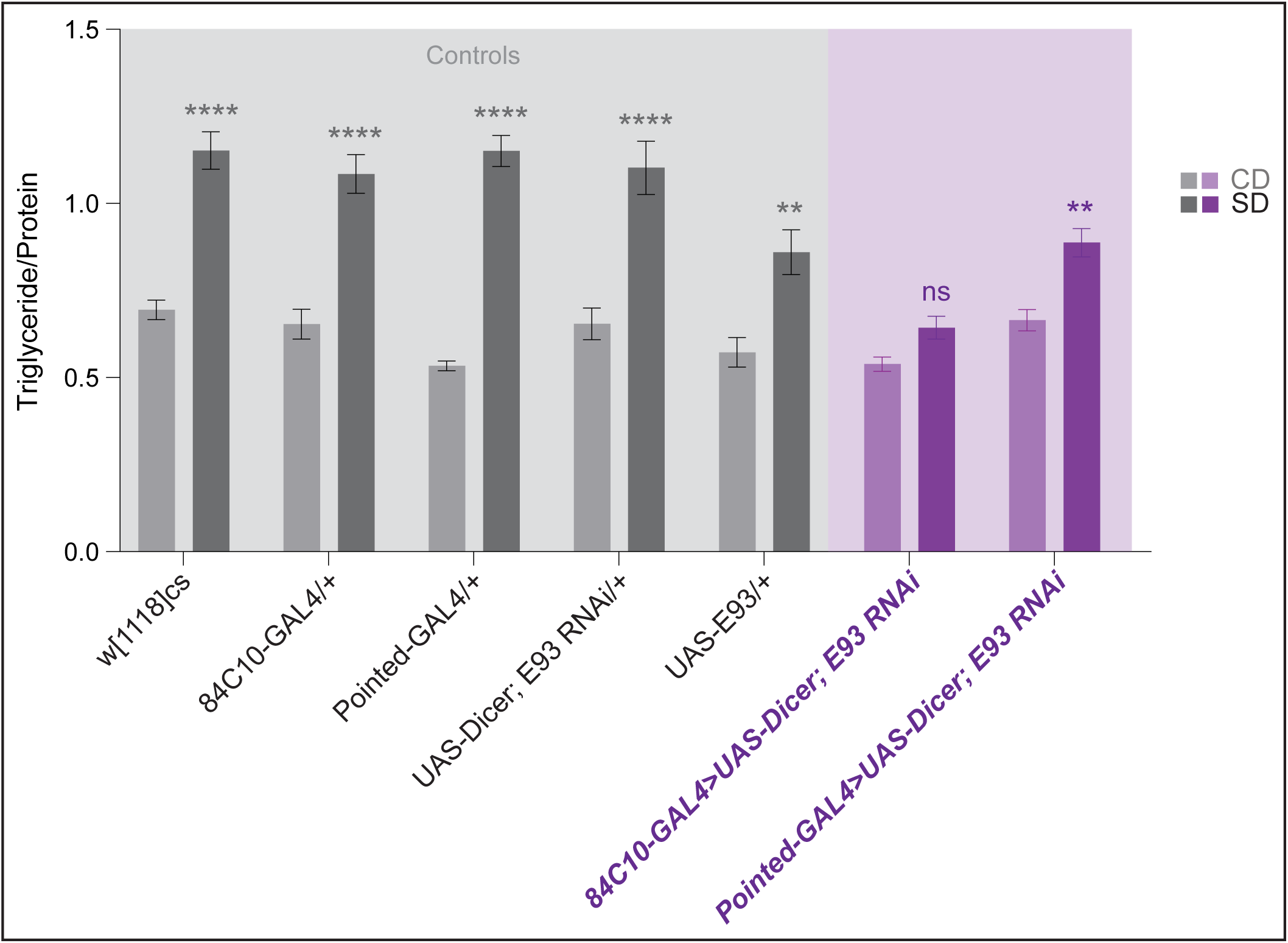
Postmitotic E93 in 84C10 dFB Neurons Is Required for Diet-Induced Triglyceride Accumulation. Triglyceride (TAG) levels normalized to total protein content in age-matched adult male flies subjected to dietary manipulation. Flies of the following genotypes were analyzed: w¹¹¹⁸ Canton-S (control), 84C10-GAL4 > w¹¹¹⁸CS, 84C10-GAL4 > UAS-Dicer; E93 RNAi, w¹¹¹⁸CS > UAS-E93, Pointed-GAL4 > w¹¹¹⁸CS, UAS-Dicer; E93 RNAi > w¹¹¹⁸CS and Pointed-GAL4 > UAS-Dicer; E93 RNAi. Flies were placed on a control or 30% sugar diet for 7 days. Data are shown as mean ± SD. *n* = 16-24 flies per condition. Statistical significance was assessed using two-way ANOVA followed by Tukey’s multiple comparisons test, with comparisons made relative to the control-diet condition. Asterisks denote significance levels as indicated.

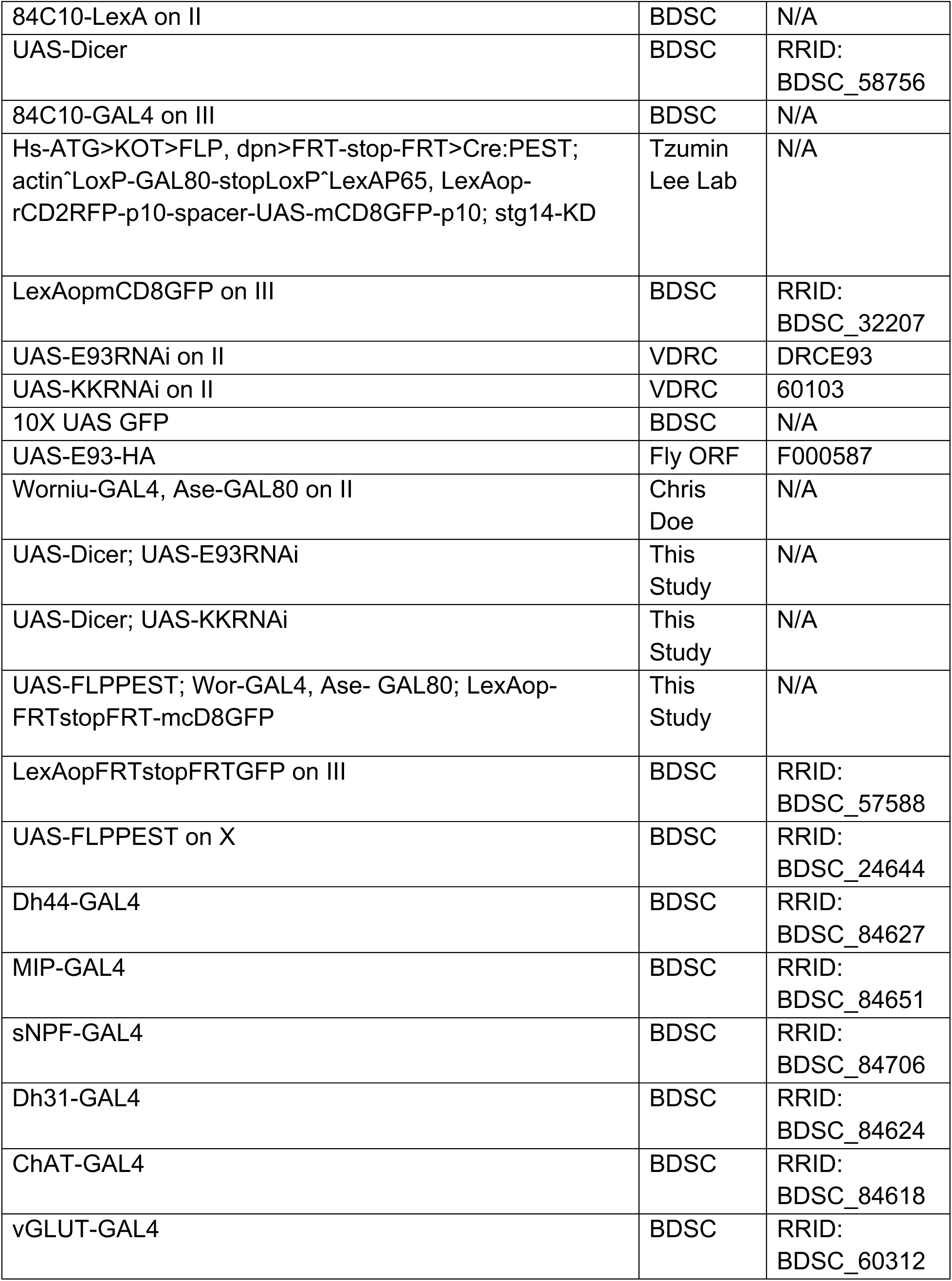

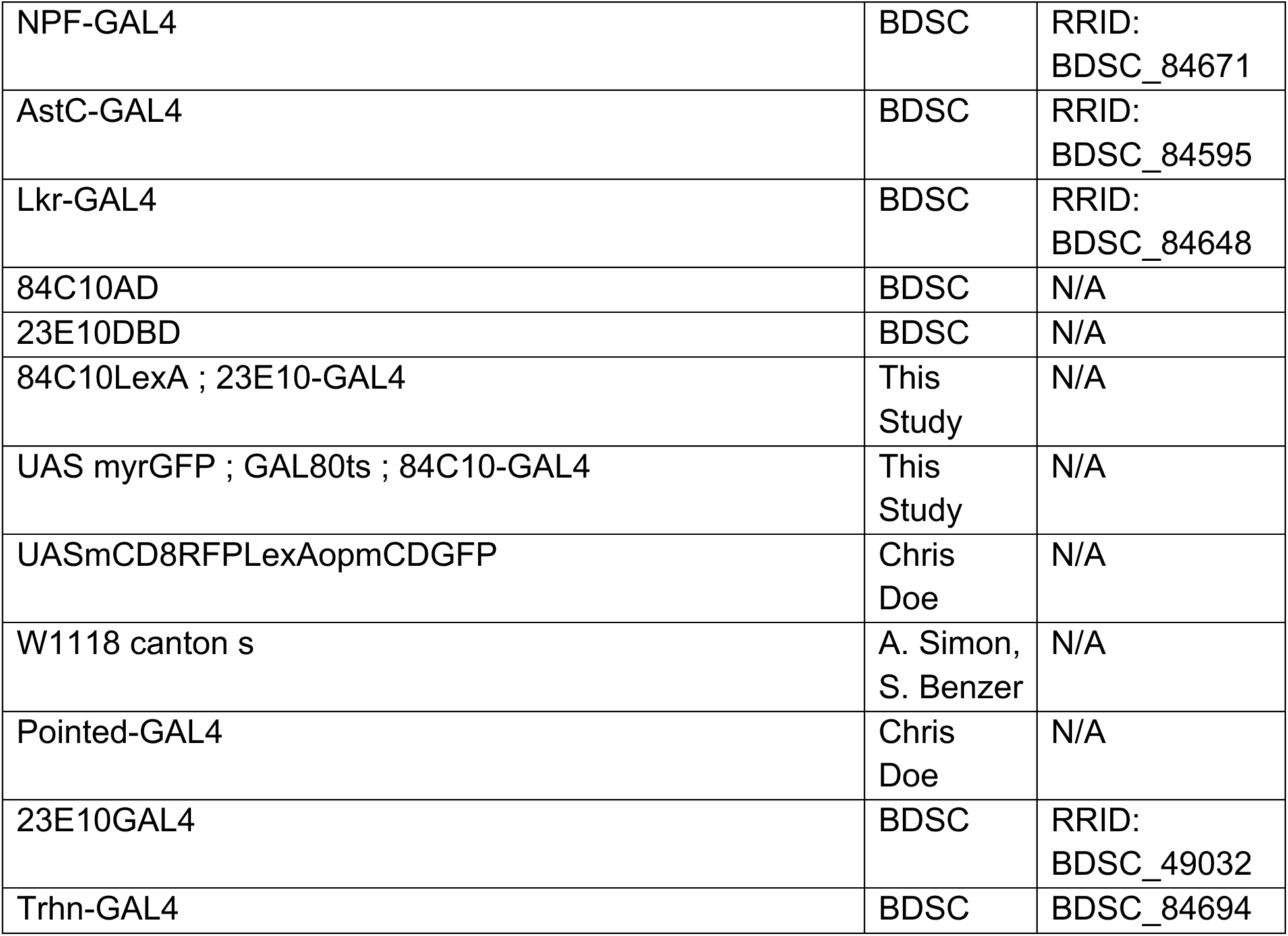

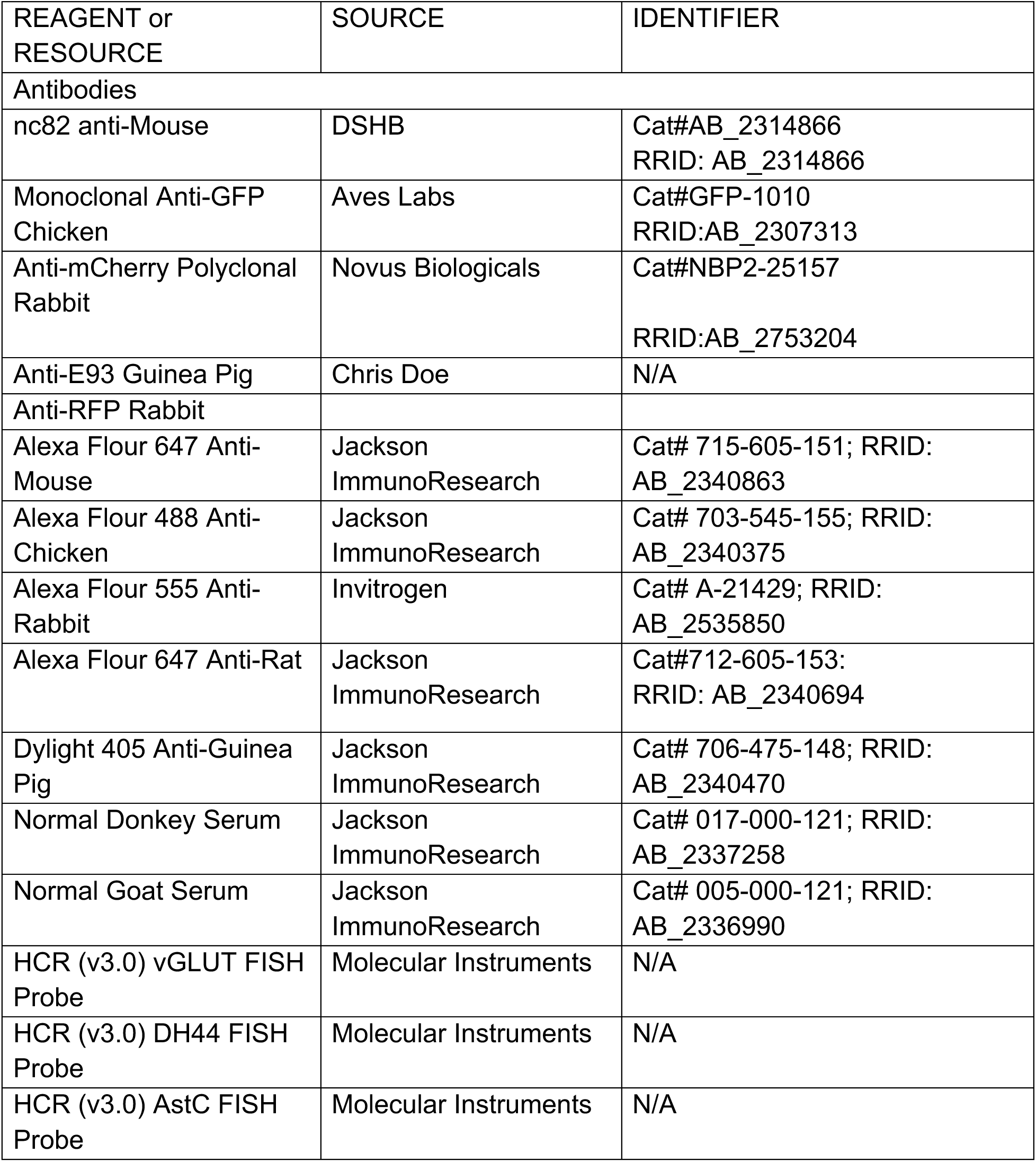

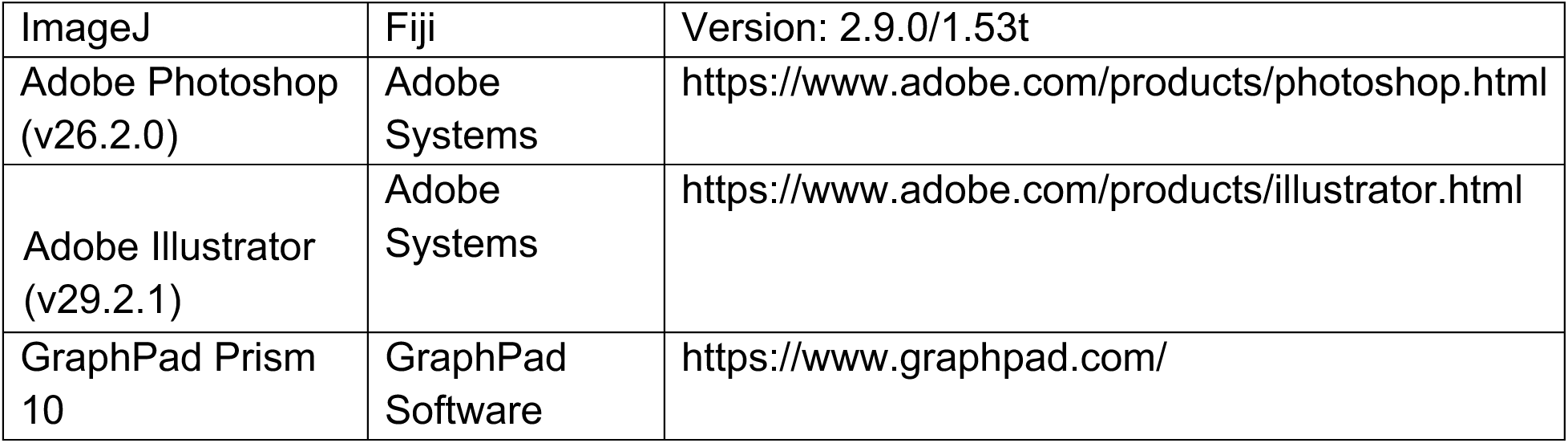

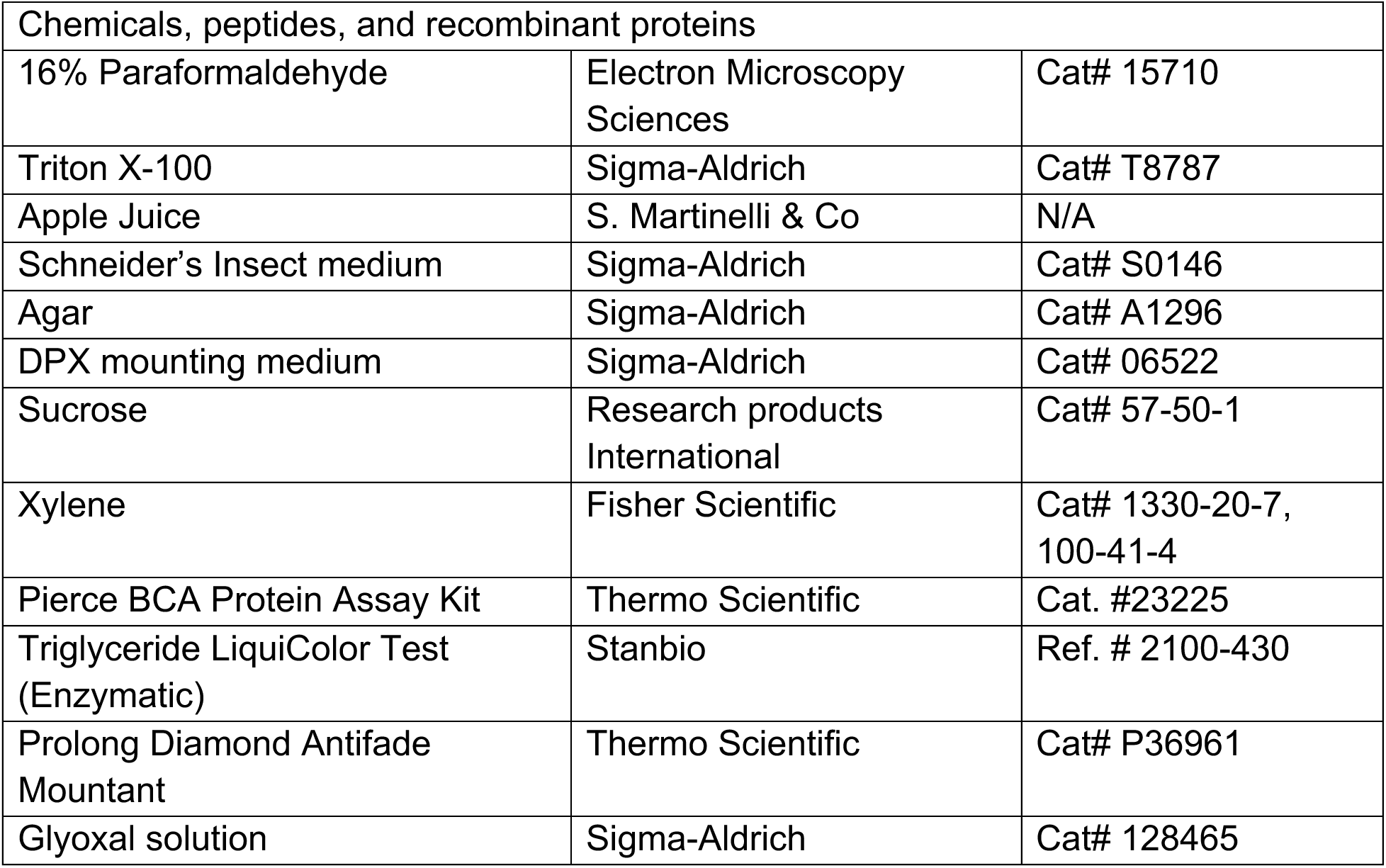

## Notes

### Competing Interest Statement

The authors have declared no competing interest.

### Summary of Updates

We have updated the text and also organized the figures to better tell our story.

